# Progressive bilateral recruitment and resilient network reorganization during temporal lobe epileptogenesis

**DOI:** 10.64898/2026.01.27.701979

**Authors:** Fabien Friscourt, Maëlle Hernot, Guru Prasad Padmasola, Clothilde Ferreira, Karl Schaller, Christoph M. Michel, Charles Quairiaux

**Affiliations:** Department of Basic Neuroscience, University of Geneva, Geneva, Switzerland; Neurosurgery Clinic, Department of Clinical Neuroscience, University Hospital Geneva, Geneva, Switzerland

**Keywords:** Temporal lobe epilepsy, Epileptogenesis, Kainate model, Hippocampal networks, Contralateral hippocampus, High-frequency oscillations, Network synchronization, Chemogenetic silencing

## Abstract

**Background:** Temporal lobe epilepsy (TLE) often originates from focal hippocampal injury but progressively evolves into a bilateral epileptic network engaging both hippocampi and distributed cortical regions. A mechanistic understanding of how this network emerges, and whether early perturbation of specific nodes can alter its trajectory, is essential for developing network-level therapeutic strategies.

**Objective:** We used a kainate-induced rodent model of TLE to (1) characterize the spatiotemporal emergence of epileptic discharges during the latent phase, (2) determine how bilaterally synchronized events develop, and (3) test whether transient chemogenetic silencing of either the ipsilateral epileptogenic focus (EF) or the contralateral hippocampus (CH) modifies large-scale epileptogenesis.

**Methods:** Freely moving mice were implanted with multi-site electrodes spanning bilateral hippocampal subfields (dentate gyrus, CA1, subiculum) and cortical regions (M2, Cg1, PrL, V1, entorhinal cortex). Longitudinal LFP recordings were performed every other day during the latent and early chronic phases following KA or saline injection. DREADD-based chemogenetic inhibition of glutamatergic neurons was applied between days 2–7 post-KA. Epileptiform events were quantified via spike rates, waveform metrics, high-frequency oscillations (HFOs), and short-latency interregional co-spiking

**Results:** Early after KA, epileptic spiking emerged locally in the ipsilateral dentate gyrus and progressively organized into HFO-coupled discharges. Contralateral hippocampal recruitment followed a distinctive biphasic time course, characterized by transient early activation, subsequent suppression, and later re-emergence with increasing bilateral coactivation. Cortical regions gradually developed higher spike rates and enhanced DG-related co-spiking, indicating large-scale network integration. Ipsilateral silencing modified local spike composition but did not prevent global network progression, whereas contralateral silencing accelerated ipsilateral epileptogenesis and strengthened pathological HFO expression.

**Conclusion:** Epileptogenesis in the KA model reflects a transition from a focal hippocampal insult to a resilient, bilateral cortico-hippocampal network. Targeting a single hippocampal node—even at early latent stages—is insufficient to halt this progression, highlighting the need for network-level therapeutic strategies.

**Graphical abstract:** 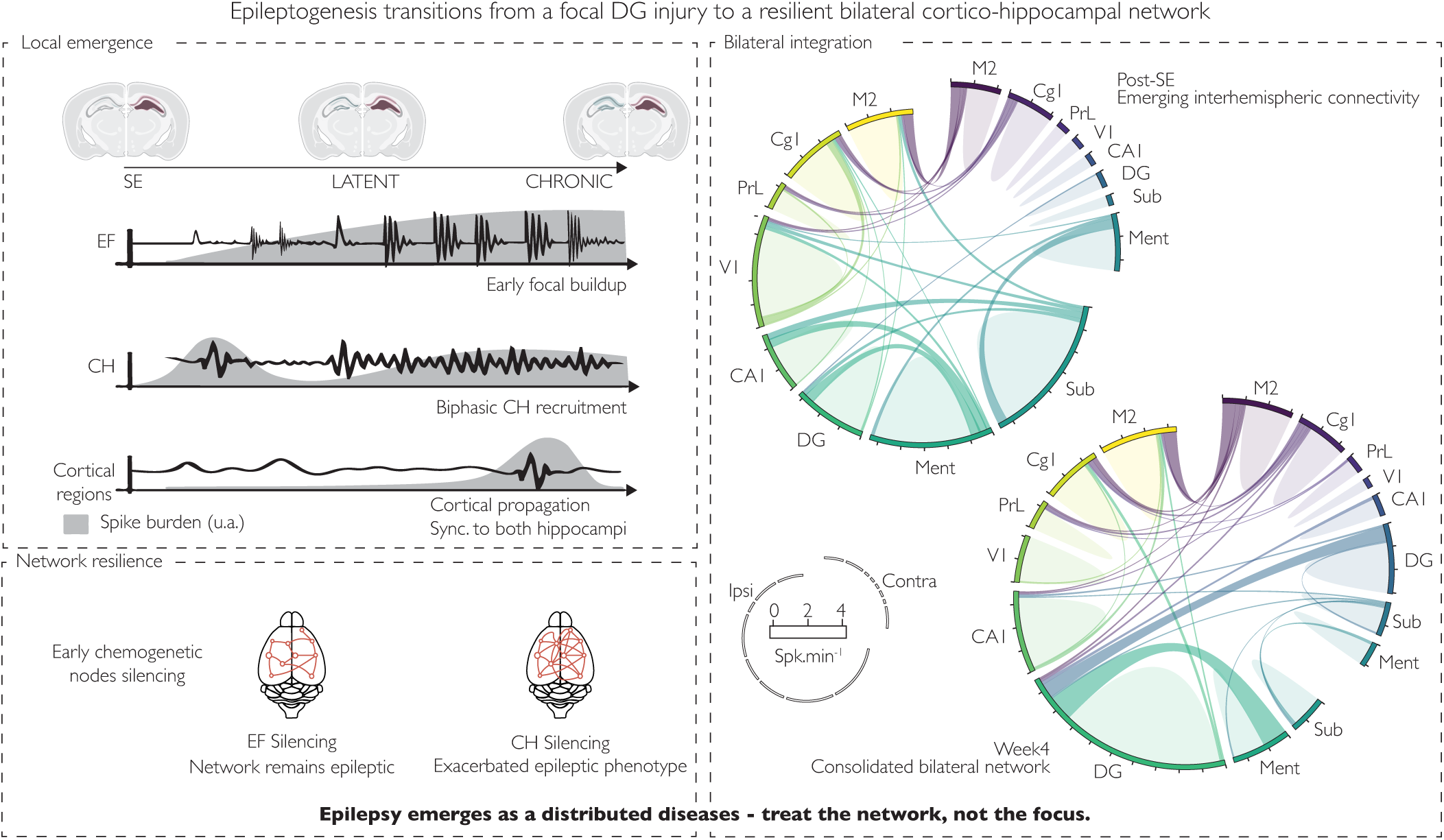

## Introduction

Temporal lobe epilepsy (TLE) is one of the most common and treatment-resistant forms of epilepsy, often characterized by seizures originating from mesial temporal structures such as the hippocampus and entorhinal cortex (Engel et al., 2013; Pitkänen & Lukasiuk, 2011). Despite advances in pharmacological treatments, a significant subset of patients with TLE remain refractory to medication, underscoring a critical need for a deeper understanding of the underlying mechanisms and novel therapeutic strategies (Goldberg & Coulter, 2013; Löscher, 2002). TLE typically develops through a “latent phase” following an initial brain insult - such as status epilepticus (SE) - during which neuronal hyperexcitability gradually increases until spontaneous seizures emerge (Curia et al., 2008; Pitkänen & Lukasiuk, 2011). Notably, approximately 20-50% of patients with TLE, particularly those with hippocampal sclerosis, have a history of SE, often in the context of febrile illness (Blumcke et al., 2017). Elucidating how neural circuits evolve and reorganize during this latent period is fundamental for early detection, intervention, and potentially disease prevention.

Among experimental models of TLE, the kainate (KA) rodent model has been extensively used due to its ability to replicate hallmark features of human mesial temporal lobe epilepsy, including hippocampal sclerosis, recurrent seizures, and a distinct latent phase (Curia et al., 2008; Lévesque & Avoli, 2013). In this model, an injection of KA (an excitotoxic glutamate analogue) into the hippocampus induces robust focal damage and triggers epileptogenic processes that gradually engage surrounding and distant brain regions, thereby mirroring the expansion of epileptiform activity seen in patients. Although the ipsilateral hippocampus (where KA is injected) is typically the first to show overt signs of hyperexcitability, studies indicate that the contralateral hippocampus (CH) and extrahippocampal regions eventually become part of the epileptic network (Bragin et al., 1999; Sheybani et al., 2018). This gradual bilateralization of hippocampal activity, sometimes accompanied by cortical involvement, reflects the inherent complexity and plasticity of epileptic circuits (Scharfman, 2007).

A growing body of evidence suggests that pathological network synchronization - measured via interictal spikes, high-frequency oscillations (HFOs), and prolonged paroxysmal discharges - may spread beyond the initially lesioned site to recruit additional structures, including neocortical and subcortical regions(Engel et al., 2013; Goldberg & Coulter, 2013; Jelerys et al., 2012; Krook-Magnuson et al., 2013; Padmasola et al., 2024; Sheybani et al., 2018, 2019). Such network-wide changes pose challenges for targeted interventions, as dampening excitability in a single locus can be offset by parallel or compensatory pathways. Recently developed chemogenetic tools, such as DREADDs (Designer Receptors Exclusively Activated by Designer Drugs), enable targeted inhibition or activation of specific neuronal populations and have shown promise in dissecting epileptic networks (Alexander et al., 2009; Urban & Roth, 2015).

In light of these persistent challenges in understanding and interrupting network-level epileptogenesis, the present study aimed to (1) characterize the spatiotemporal evolution of epileptic discharges with high spatial and temporal resolution across hippocampal and extra-hippocampal structures, (2) identify the emergence of large-scale network synchrony during the latent phase, and (3) determine whether transient inhibition of either the ipsilateral or contralateral hippocampal node during the early latent phase could attenuate the development of epileptiform activity within the focus and the broader network. Because epileptogenic circuits are highly plastic and rapidly reorganize, we hypothesized that silencing a single hippocampal node would be insufficient to prevent large-scale epileptogenesis, as compensatory strengthening of interhemispheric and cortico-hippocampal pathways would maintain pathological synchronization. By combining multi-electrode recordings, LFP recordings analyses, and targeted DREADD inhibition, we provide new insights into how focal lesions evolve into widely distributed epileptogenic networks and discuss the implications for future therapeutic interventions in TLE.

## Materials and Methods

### Animals

Thirty-five male C57BL/6J mice (Charles River Laboratories), aged 12–15 weeks, were randomly separated to four groups: (i) KA-injected (Epileptic; *n* = 12), (ii) saline-injected controls (Saline; *n* = 5), and two cohorts receiving a pre-KA viral injection targeting either (iii) the future epileptic focus (EF silencing; *n* = 9) or (iv) the contralateral hippocampus (CH silencing; *n* = 9). Before KA, mice were group-housed on a 12 h light–dark cycle (lights on 07:00–19:00); after KA they were singly housed under identical conditions. All experiments were conducted during the light phase. Experimental timelines are shown in Fig. 1A. All procedures complied with Swiss National Guidelines for Animal Experimentation and were approved by the Geneva Cantonal Veterinary Office (GE190).

**Figure 1:**
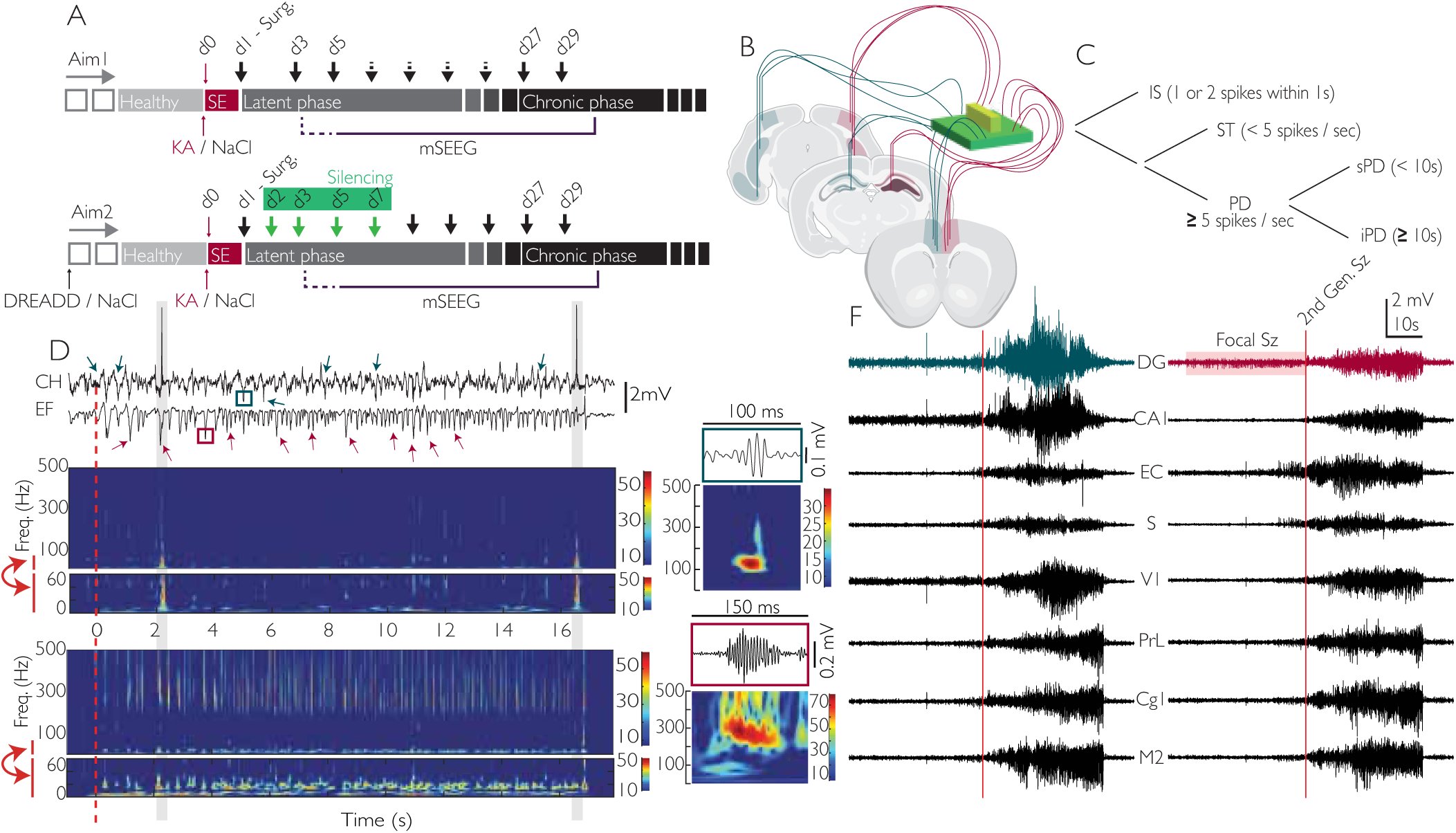
Experimental design and classification of epileptic events across epileptogenesis. A | Experimental timeline for the two study aims. Aim 1 (top): characterization of epileptogenesis following kainate (KA) or saline injection at day 0 (red). Animals underwent electrode implantation (Surg.) and were monitored during the latent (gray) and chronic (black) phases. Electrophysiological recordings (black arrows) were performed every other day (d3–d25) and at later chronic time points (d27–d29). Aim 2 (bottom): chemogenetic network modulation. In addition to KA, animals received inhibitory DREADDs or saline injection, with chemogenetic silencing (green) applied during the (d2-d7). Multi-site intracranial EEG (mSEEG) recordings were performed throughout the latent and chronic phases. B | Schematic representation of electrode implantation in freely moving mice. Six multichannel electrode bundles were stereotaxically implanted in bilateral hippocampal and cortical regions and connected to an external electrode interface board (EIB). Targeted areas included hippocampal subfields (dentate gyrus, CA1, subiculum) as well as cortical regions (entorhinal cortex, prelimbic cortex, cingulate cortex, secondary motor cortex, and primary visual cortex). The schematic illustrates the spatial distribution of electrodes across both hemispheres. Exact stereotaxic coordinates and surgical procedures are provided in the Materials and Methods. C | Schematic classification of epileptiform events based on spike rate and duration. Events were first categorized as isolated spikes (IS, one or two spikes occurring within 1 s). When multiple spikes were detected, events were further classified as spike trains (ST, <5 spikes·s^-1^) or paroxysmal discharges (PD, ≥5 spikes·s^-1^). PDs were then subdivided according to their duration into short PDs (sPD, <10 s) and intermediate/long PDs (iPD, ≥10 s). Full classification criteria are provided in the Materials and Methods. D | Representative example of a paroxysmal discharge recorded at day 27 post-KA. Top: raw traces from the contralateral hippocampus (CH) and the epileptogenic focus (EF). Arrows highlight examples of typical IS in each region. The grey rectangle illustrated a network epileptic discharge. Bottom: corresponding time–frequency (TF) decompositions. The EF trace displays a high-load epileptic event characterized by prominent spike bursts in the 4–40 Hz band, accompanied by robust high-frequency oscillations (HFOs), particularly in the fast-ripple range (250–500 Hz). In the CH, simultaneous but less intense activity is observed, with spikes in the 4–40 Hz band associated with HFOs predominantly in the ripple range (80–250 Hz). Insets on the right illustrate representative examples of ripple (top) and fast-ripple (bottom) events with their respective TF signatures. E | Representative example of a secondarily generalized seizure recorded in a chronically epileptic mouse. Local field potentials (4–40 Hz band) are shown simultaneously across multiple brain regions, including hippocampal subfields (dentate gyrus, DG; CA1), entorhinal cortex (EC), subiculum (S), primary visual cortex (V1), prelimbic cortex (PrL), cingulate cortex (Cg1), and secondary motor cortex (M2). The red vertical lines indicate the onset of the focal seizure in the DG (left) and the subsequent secondary generalization (right). Initially, the discharge remains confined to the DG, but as the event evolves it rapidly recruits hippocampal and neocortical regions, illustrating the transition from a focal to a widespread, highly synchronized network state. The upper schematic indicates electrode placements in hippocampal and cortical regions.

### Chemogenetic Manipulations

To silence principal neurons, an AAV carrying hM4Di under the CamKIIα promoter (rAAV1-CamKII2-hM4Di-mCherry; UNC Vector Core) or saline was delivered stereotaxically to dorsal and ventral hippocampus of 10–12-week-old mice under isoflurane anesthesia. Coordinates (relative to Bregma) were dorsal: AP −2.2 mm, ML ±1.0 mm, DV 1.7 mm; ventral: AP −3.3 mm, ML ±2.0 mm, DV 1.9 mm. One microliter was injected per site at 10 nL·s^-1^ via a pulled glass pipette. KA induction was performed three weeks after viral injection. To activate DREADDs, clozapine-N-oxide (CNO; 2.5 mg/100 mL in 4% sucrose) was administered in the drinking water from day 2 to day 7 post-KA. Further device details and consumables are listed in Supplementary Methods.

### Kainate injection (epilepsy induction)

TLE was induced by a status epilepticus (SE) caused by a unilateral stereotaxic injection of kainic acid (0.35 nmol, 5 mM, 70 nL; 10 nL·s^-1^; Tocris) into the left dorsal hippocampus (AP −1.8 mm, ML −1.6 mm, DV 1.9 mm). The injection capillary remained in place for 3–4 min to permit diffusion before closure. Saline controls received 0.9% NaCl. Subsequent procedures (e.g., electrode implantation) were performed ≥24 h later (day 0 = KA). Full surgical steps are detailed in Supplementary Methods.

### Multi-region electrode implantation

At day 1 post-KA, mice were anesthetized (medetomidine, midazolam, fentanyl) and implanted with 16 PFA-coated stainless-steel electrodes arranged in six polytrodes targeting eight bilateral regions: secondary motor cortex (M2), cingulate (Cg1), prelimbic (PrL), primary visual cortex (V1), hippocampal CA1, dentate gyrus (DG), subiculum (Sub), and entorhinal cortex (Ment) (Fig. 1B). Electrodes were gold-plated to ∼400 kΩ and fixed on the skull with UV-curable resin and dental cement; reference and ground screws were placed over the cerebellum. Post-operative care included analgesia, antibiotics, and nutritional gel. Full stereotaxic coordinates, implant geometry, materials, and fabrication steps are in Supplementary Methods.

### Freely moving recordings

From day 3 to day 29 post-KA, neural activity was recorded every other day (∼1 h/session) using a Digital Lynx SX system (Neuralynx; 32 kHz; 0.1–9000 Hz bandwidth) to capture LFPs and multi-unit activity. Sessions were performed in the afternoon to minimize circadian variability. To reduce handling stress, animals were briefly anesthetized for tethering; recordings commenced ≥10 min after full locomotor recovery. Additional handling details are provided in Supplementary Methods.

### Epileptiform spike detection

A semi-automated MATLAB pipeline was used adapted from Heining et al., 2019(Heining et al., 2019). Signals were downsampled to 2 kHz, low-pass filtered at 1 kHz, and artifacted channels excluded (Fig. 2A). In the 4–40 Hz band, spikes were defined as events exceeding baseline by 5 SD, with an 80 ms refractory period. To avoid duplicates, spikes detected within 5 ms across electrodes of the same polytrode were cleaned. Comprehensive preprocessing parameters, artifact rejection rules, and quality controls are reported in Supplementary Methods.

**Figure 2:**
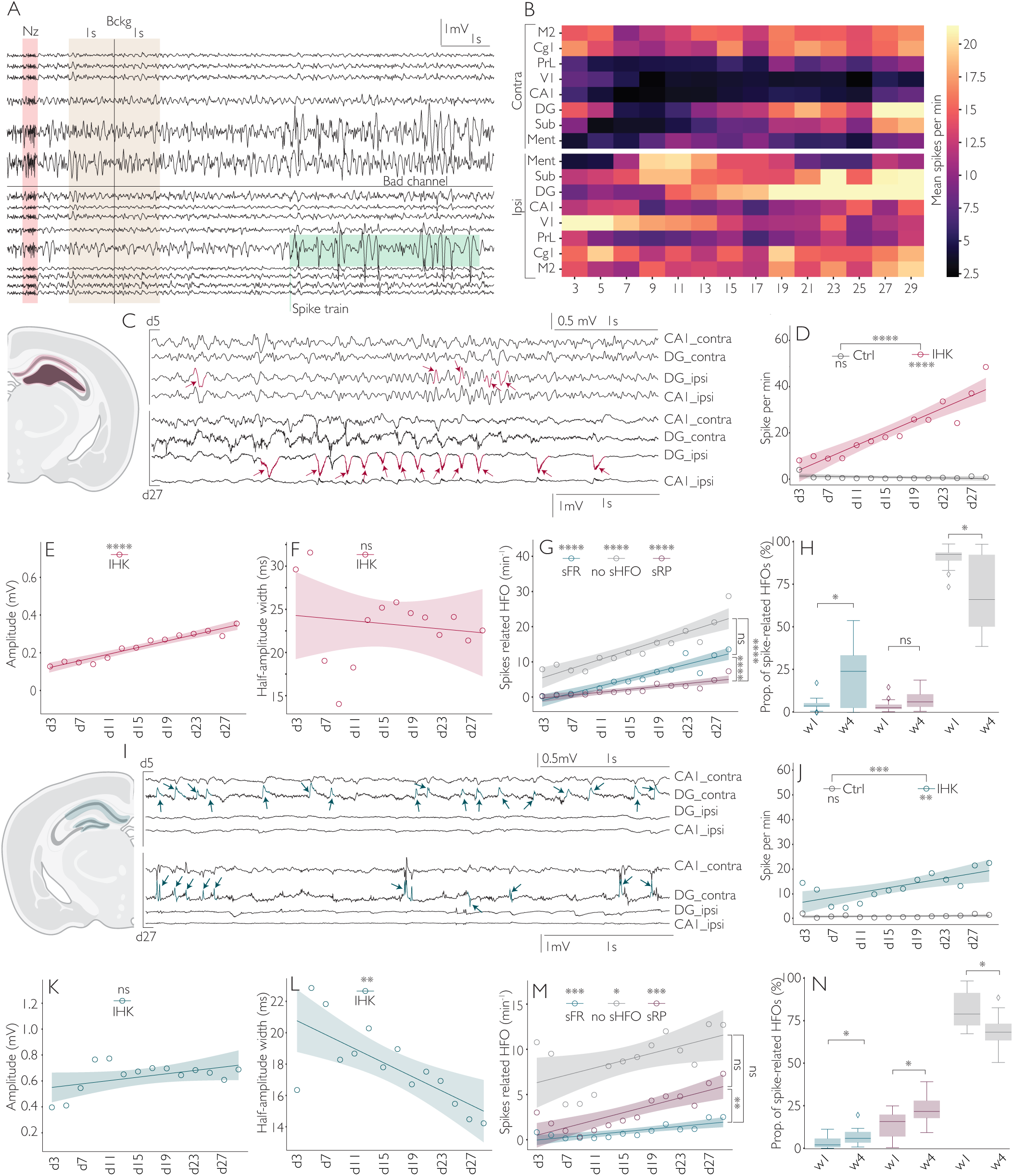
Progressive emergence and organization of epileptiform activity in both hippocampi. A | Example trace & analysis workflow. Raw LFP after denoising (Nz) (notch/line-noise + artifact removal). A manually placed background (Bckg) marker indicates 2-s windows free of epileptiform activity used as baseline. Bad channels are flagged and excluded. Epileptic events, such as spike trains (green) are then automatically detected on the remaining channels. B | Heatmap of mean spikes per minute for each electrode (rows) across recording days (columns) in KA-treated mice (IHK). Rates were computed per (mouse, day, electrode, recording), normalized by minutes, and averaged across mice for each (day, electrode). Color scale capped at the 95th percentile to limit outlier influence. C | Representative hippocampal raw traces (ipsilateral to the KA injection). Focal spiking recorded in EF at day 5 (top) and day 27 (bottom) post-KA. Simultaneous CA1 and contralateral DG traces illustrate the later emergence of large-amplitude, recurrent spikes. D | Mean spike rate over time (ipsilateral DG) with best-fit lines (OLS). Daily spike rate in epileptic (red, n=12) vs control (gray, n=5) animals. Ordinary least-squares linear regression (OLS) fits with mouse-clustered SEs; shaded bands are 95% CIs. Epileptic mice show a robust increase (slope = 1.3640, R² = 0.8655, p < 0.0001); controls are flat (slope = –0.0421, R² = 0.1297, p = 0.2059). Pairwise slope test: Ctrl vs IHK, p< 0.0001. E | Spike amplitude (ipsilateral). Mean amplitude increases across days (slope = 8.6305 µV·day^-1^, R² = 0.9363, p < 0.0001; shaded = 95% CI). F | Half-amplitude width (ipsilateral). Largely stable over time (slope = –0.077 ms·day^-1^, R² = 0.021, p = 0.62; shaded = 95% CI). G | Spike-related HFOs per minute (ipsilateral). Rates for spikes with fast ripples (sFR, blue), ripples (sRP, red), and no HFO (noHFO, grey) (OLS + clustered SEs; shaded = 95% CI): sFR slope = 0.5114, R² = 0.8915, p < 0.0001; sRP slope = 0.2042, R² = 0.7622, p < 0.0001; noHFO slope = 0.6476, R² = 0.8197, p < 0.0001. Pairwise slope tests: sFR vs sRP, p < 0.0001; sRP vs noHFO, p < 0.0001; sFR vs noHFO, p = 0.1802. H | Fraction of spike-related HFOs (ipsilateral hippocampus). Same color code as in panel G. Each dot = mouse-level proportion; boxes summarize inter-mouse distribution (median + q25-75 + whisker max/min values). Paired Wilcoxon between week4 post-K injection (W4, d23 → d29) vs week1 post-K injection (W1, d3 → d7) with BH–FDR within panel: sFR W1 = 0.0513 → W4 = 0.2262, p = 0.0323; sRP W1 = 0.0441 → W4 = 0.0750, p = 0.11; noHFO W1 = 0.9047 → W4 = 0.6989, p = 0.0171. I | Representative raw traces in the hippocampus contralateral to the KA injection (CH). Focal spiking in CH at day 5 (top) and day 27 (bottom); simultaneous CA1 and ipsilateral DG for comparison. J | Spike rate over time (contralateral). Daily spike rate in epileptic (blue, n=12) vs control (gray, n=5) animals (OLS + clustered SEs; 95% CI). Epileptic mice show a later, smaller increase (slope = 0.4898, R² = 0.5313, p = 0.0031); controls remain stable (slope = 0.0152, R² = 0.0533, p = 0.4270). Pairwise slope test: Ctrl vs IHK, p <0.001. K | Spike amplitude (contralateral). Trend toward increase (slope = 6.5876 µV·day^-1^, R² = 0.2348, p = 0.0791; 95% CI). L | Half-amplitude width (contralateral). Significant narrowing over time (slope = –0.2217 ms·day^-1^, R² = 0.535, p = 0.00293; 95% CI). M | Spike-related HFOs per minute (contralateral). sFR slope = 0.0759, R² = 0.6388, p = 0.0006; sRP slope = 0.2070, R² = 0.6842, p = 0.0003; noHFO slope = 0.2026, R² = 0.3330, p = 0.0003. Pairwise slope tests: sFR vs sRP, p = 0.0028; sFR vs noHFO, p = 0.1334; sRP vs noHFO, p = 0.9612. N | Fraction of spike-related HFOs (contralateral). Paired Wilcoxon (W4 vs W1) with BH–FDR within panel: sFR W1 = 0.0366 → W4 = 0.0729, p = 0.0479; sRP W1 = 0.1466 → W4 = 0.2291, p = 0.040; noHFO W1 = 0.8168 → W4 = 0.6980, p = 0.0171.

### Classification of epileptiform discharges

Epileptiform activity was categorized (Fig. 1C) as: (i) isolated spikes (IS), separated by ≥1 s; (ii) spike trains (ST), with interspike intervals <1 s but never reaching 5 spikes·s^-1^; (iii) paroxysmal discharges (PD), reaching ≥5 spikes·s^-1^ for >1 s; and (iv) in addition to PDs, we defined ictal paroxysmal discharges (iPDs) as PDs lasting ≥10s, consistent with prior work in the KA model (Padmasola et al., 2024). All iPDs were visually reviewed by two independent, blinded raters (MH, CQ) to confirm seizures. Network-level interictal events spanning >5 bundles within 100 ms were excluded from single-region analyses and categorized as network IED. Event grouping across electrodes and rules for contralateral coincidences are expanded in Supplementary Methods. In the following analyses, the epileptogenic focus (EF) refers to the ipsilateral dentate gyrus, corresponding to the site of KA injection and the earliest appearance of epileptiform discharges.

### High-Frequency Oscillations (HFOs)

HFOs associated with spike peaks were classified as ripples (RP, 80–250 Hz) or fast ripples (FR, 250–500 Hz) within a ±40 ms window. Events crossing amplitude thresholds for a minimum number of cycles were labeled accordingly (Fig. 1D). Exact thresholds, cycle criteria, duration estimation, and frequency derivation are detailed in Supplementary Methods.

### Histology

At day 29, mice were deeply anesthetized and perfused with 0.9% NaCl followed by 4% paraformaldehyde. Brains were post-fixed, sectioned at 80 µm, and native mCherry fluorescence was used to verify DREADD expression under identical exposure settings across animals. Additional staining (e.g., DAPI), imaging hardware, and acquisition parameters are provided in Supplementary Methods.

### Statistical analyses

Throughout the figures, longitudinal trends are displayed using mean values with linear regression fits, whereas distributional comparisons use boxplots showing median and interquartile ranges. In addition, we applied supervised machine-learning classifiers (leave-one-subject-out cross-validation) to evaluate whether early electrophysiological features could predict later seizure severity (Pedregosa et al., 2011). The full statistical framework - including computation of spike-rate heatmaps, regression models for longitudinal trends, seizure-latency analysis, spectral power metrics, and additional distributional and network-sync analyses - is described in the Supplementary Materials and Methods with references to the corresponding Supplementary Figures.

## Results

### Early focal spike emergence and HFO maturation

From the first recording day after KA injection (d3), epileptic mice displayed sporadic interictal spikes that progressively increased in frequency and temporal organization during the latent phase (Fig. 2C). Spatial mapping of spike rates (Fig. 2B) revealed a stereotyped recruitment sequence: activity first emerged in the EF, then spread to the ipsilateral subiculum (Sub_ipsi), and later engaged contralateral hippocampal and cortical regions. A subset of animals showed transient early spiking in V1 - a cortical activity pattern likely reflecting acute post-SE hyperexcitability - which rapidly subsided, giving way to the more canonical hippocampal-first recruitment. In contrast, saline-treated controls remained largely silent across the entire recording period.

Within the epileptic hippocampus, spike rates increased steadily across days (Fig. 2D), accompanied by a gradual rise in spike amplitude (Fig. 2E) while spike width remained stable (Fig. 2F), indicating enhanced synchronous recruitment of local neurons in the spiking events. High-frequency oscillations (HFOs) provided further insight into the maturation of epileptiform activity: both ripple-related (sRP) and fast-ripple–related (sFR) spikes increased over time (Fig. 2G), but the relative increase was significantly greater for sFR, whereas the proportional rise in sRP did not reach significance. Notably, spikes uncoupled from HFOs remained the most prevalent subtype throughout the latent phase, although their proportion decreased by week 4 (Fig. 2H). By week 4, sFR constituted a significantly larger fraction of HFO-coupled discharges than sRP (Fig. 2G–H).

Altogether, these results suggest that epileptogenesis is not a uniform amplification of excitability but a structured synchronization process. The increasing prominence of sFR over sRP suggests a transition from diffuse early hyperexcitability toward the emergence of more locally resonant, hyperconnected microcircuits within the ipsilateral hippocampus, marked by increasingly synchronized, HFO-coupled events.

### Delayed contralateral recruitment with biphasic progression

Contralateral hippocampal recruitment followed a biphasic time course. In the days immediately following SE, contralateral spiking was detectable, but it transiently decreased around days 7–9 before rising again toward the chronic stage (Fig. 2I–J). This pattern indicates that contralateral activation is initially weak and unstable but strengthens later as interhemispheric coupling matures.

Contralateral spike morphology evolved in parallel with ipsilateral changes, with spike amplitude increasing slowly and spike width decreasing significantly over time (Fig. 2K–L). Similarly to the ipsilateral side, contralateral spikes also became increasingly coupled to HFOs, although ripples predominated, with fast ripples, even if increasing with time, remaining comparatively rare (Fig. 2M–N). By week 4, sRP represented a significantly larger fraction of contralateral events than during week 1, highlighting a hemispheric asymmetry in the maturation of epileptiform activity, whereas non-HFO spikes declined.

Together, these observations indicate that contralateral circuits progress along a slower and less pathological maturation trajectory than the ipsilateral side. Yet, the biphasic recruitment, the evolving spike morphology, and increasing ripple coupling demonstrate that they still shift from an initially unstable state to a more organized epileptiform regime by the chronic stage.

### Spectral reorganization reveals hemispheric divergence

To assess how epileptogenesis alters background network dynamics, we quantified longitudinal power spectral density (PSD) in both hippocampi. In the CH, PSD remained largely stable across weeks (Fig. 3A, C–D, G–H). Week-wise comparisons of PSD at W1 and W4 (Fig. 3A) revealed only minor, mostly non-significant temporal fluctuations, except for a transient decrease in delta power at W1 as compared to controls. Consistently, linear day-wise models (Fig. 3 C-D, G-H) showed shallow, non-significant slopes for delta, theta, beta, and gamma relative power, with no interaction with cohort, indicating preservation of spectral composition in CH despite ongoing epileptogenesis.

**Figure 3:**
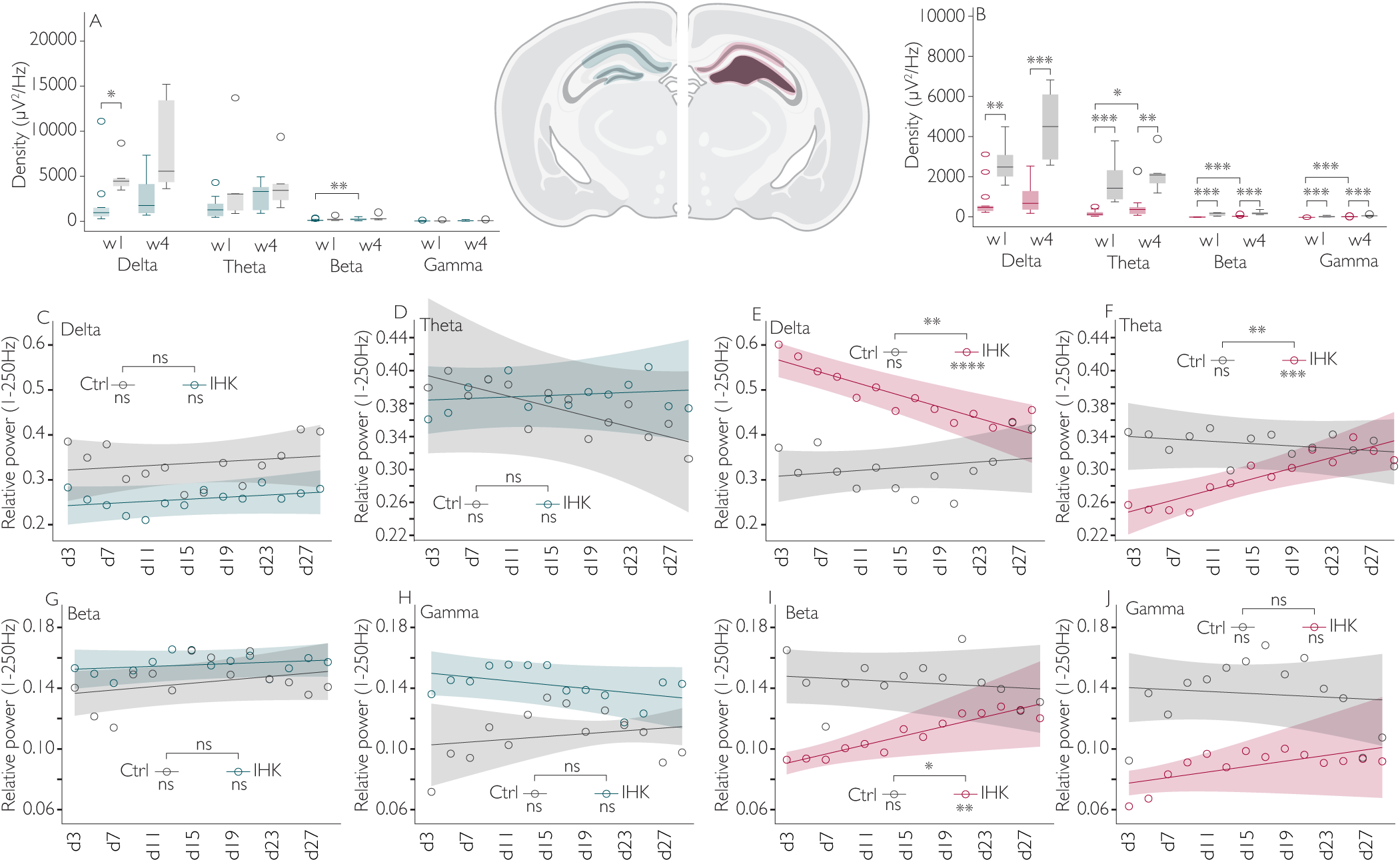
Ipsilateral spectral remodeling contrasts with contralateral stability. A | Weekly PSD density (µV²·Hz^-1^) in CH. Week-wise boxplots at W1 and W4 for delta, theta, beta, and gamma in saline controls (grey) and epileptic animals (blue). Boxplots show the per-mouse distributions (median, interquartile range; whiskers = 1.5×IQR; circles = individual outliers). Contralateral distributions show only small, mostly non-significant shifts across weeks; the notable exception is delta at W1 (q = 0.0177, r_rb = −0.85). B | Weekly PSD density (µV²·Hz^-1^) in the injected hippocampus (saline, grey, KA, red). EF. Clear separation from controls emerges by W1 and widens by W4: W1: delta↓ (q=0.009, r_rb=−0.82), theta↑ (q=0.0009, r_rb=−1), beta↑ (q=0.00047, r_rb=−1), gamma↑ (q=0.00047, r_rb=−1). W4: delta↓ (q=0.0009, r_rb=−1), theta↑ (q=0.003, r_rb=−0.88), beta↑ (q=0.00062, r_rb=−0.97), gamma↑ (q=0.00047, r_rb=−0.97). (Ripple band not displayed—did not survive global FDR across weeks/electrodes in the final set.) C | CH - Delta relative power (1–250 Hz) vs day. Shallow, non-significant drift (slope=+0.00115·day^-1^, R²=0.0102, p=0.311); group×day vs Ctrl: ns. (p=0.982). D | CH - Theta relative power, small, non-significant change (p=0.727); interaction vs Ctrl: ns. (p=0.179). G | CH – Beta, small, non-significant change (p=0.418); interaction vs Ctrl: ns. (p=0.49). H | CH – Gamma, small, non-significant change (p=0.129); interaction vs Ctrl: ns. (p=0.155). E | EF – Delta, monotonic decline (slope=−0.0063·day^-1^, R²=0.204, p=2.49×10⁻⁷); group×day interaction significant vs Ctrl (p=0.0003). F | EF – Theta, increase over days (slope=+0.00333·day^-1^, R²=0.18, p=0.0007); interaction vs Ctrl (p=0.003). I | EF – Beta, increase over days (slope=+0.00149·day^-1^, R²=0.118, p=0.0087); interaction vs Ctrl (p=0.014). J | EF – Gamma, modest, non-significant upward trend (slope=+0.0009·day^-1^, R²=0.0302, p=0.156); interaction vs Ctrl ns. (p=0.187). Within-electrode W1→W4 (EF, BH-FDR per band). Delta_rel decreased (W=2, q=0.0029, r=−0.94); Theta_rel increased (W=7, q=0.019, r=0.79); Beta_rel increased (W=4, q=0.0068, r=0.87); Gamma_rel increased (W=11, q=0.027, r=0.685). No contralateral week-to-week changes survived correction. The EF spectrum shows a structured redistribution—delta↓, theta/beta↑ (±gamma↑)—that unfolds across days and tends toward control-like proportions by ∼d27, whereas contralateral PSD remains comparatively stable.

In contrast, the kainate-injected hippocampus exhibited significantly reduced power across all frequency bands compared to saline controls already during W1 (Fig. 3B), with these differences further accentuated by W4. Longitudinal day-wise modeling (Fig. 3E-F, I-J) revealed significant monotonic trends over time: delta relative power, initially dominant following SE, gradually declined, whereas theta and beta steadily increased, and gamma showed a modest upward trajectory. As a result, the relative contributions of frequency bands gradually shifted toward control-like values by the end of the monitoring period, suggesting partial re-equilibration of background oscillatory activity rather than continuous deterioration.

Within-electrode paired W1→W4 comparisons further supported longitudinal remodeling in EF. Delta relative power decreased across weeks, while theta, beta, and gamma increased (Fig. 3 B), indicating a redistribution of spectral weight from slow-wave to faster rhythms. No contralateral week-to-week shifts survived multiple-comparison correction, reinforcing hemispheric asymmetry.

Together, these findings demonstrate that chronic epileptogenesis induces a gradual, frequency-specific reorganization of ongoing background activity within the epileptic hippocampus. Notably, this altered spectral state evolves toward control-like proportions over time, while the contralateral hemisphere remains comparatively stable.

### Seizure onset symmetry but ipsilateral consolidation

We next examined in the bi-hippocampal network whether ictal paroxysmal discharge (iPD), which are usually considered reminiscent of seizures in the model, emerged earlier in the epileptic hippocampus than in the contralateral one. Kaplan–Meier survival analysis revealed overlapping seizure-free probabilities for EF and CH, indicating that the first iPD occurred with similar latency in both hemispheres (Fig. 4A). Despite the unilateral insult, early ictogenesis is therefore bilaterally symmetric.

**Figure 4:**
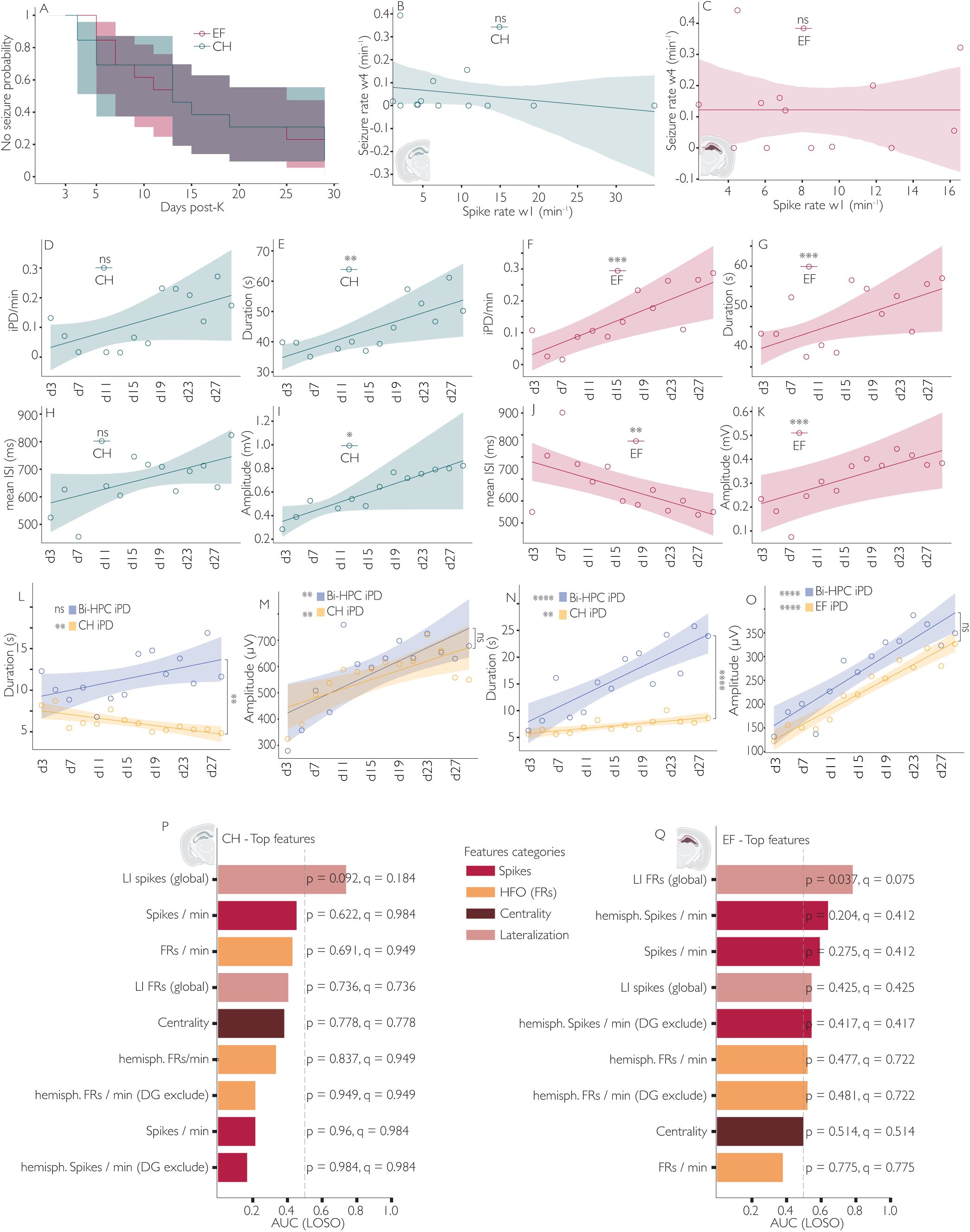
Ipsilateral amplification of paroxysmal dynamics despite symmetric seizure onset. A | Kaplan–Meier survival curves showing the latency to the first ictal paroxysmal discharge (iPD) for each electrode. Curves correspond to the ipsilateral dentate gyrus (EF, pink) and contralateral dentate gyrus (CH, blue). The y-axis represents the probability of remaining seizure-free, and the x-axis shows days post-KA. Shaded areas indicate 95% confidence intervals, and tick marks denote censored observations (animals without seizures up to their last recording). Stepwise drops indicate first-seizure events. Comparison using the log-rank test (χ² = 0.018, p = 0.894; −log₂ p = 0.162) showed no significant difference between EF and CH, indicating symmetric seizure onset latency. B–C | *Early interictal spiking does not predict later seizure rate.* Scatter plots show correlations between week1 interictal spike rate (x-axis) and week4 seizure rate (y-axis) in CH (B, blue) and EF (C, pink). Each dot represents one animal; shaded areas show 95% confidence intervals for the regression line. No significant correlation was found (CH: Spearman ρ = –0.52, p = 0.070; EF: ρ = 0.02, p = 0.957), indicating that early interictal spiking does not forecast later seizure burden. D–K | Progressive evolution of dentate iPDs after KA injection. Time courses (days 3–27) of iPD features in the contralateral hippocampus (CH; blue, left) and epileptogenic focus (EF; pink, right). Points show daily means ± SEM; shaded bands show 95% confidence intervals for the linear regression (OLS fit with mouse-clustered standard errors). CH: (D) iPD rate tended to increase (slope = +0.0068 min⁻¹·day⁻¹, R² = 0.12, p = 0.092); (E) mean duration increased significantly (+0.73 s·day⁻¹, R² = 0.19, p = 0.0084); (H) inter-spike interval (ISI) showed a non-significant upward trend (+6.45 ms·day⁻¹, R² = 0.13, p = 0.086); (I) amplitude tended to increase (+19.8 µV·day⁻¹, R² = 0.24, p = 0.053). EF: (F) iPD rate increased markedly (+0.0095 min⁻¹·day⁻¹, R² = 0.16, p = 3.0 × 10⁻⁴); (G) mean duration increased (+0.62 s·day⁻¹, R² = 0.10, p = 0.026); (J) ISI shortened significantly (–9.97 ms·day⁻¹, R² = 0.18, p = 2.0 × 10⁻⁴); (K) amplitude increased progressively (+9.24 µV·day⁻¹, R² = 0.11, p = 0.011). L - O | Regional and bilateral hippocampal PDs differ in duration and amplitude over time. (K) CH duration: regionally isolated events decreased slightly (slope = –0.108, R² = 0.54, p = 0.0027), while Bi-HPC events trended downward (slope = –0.172, R² = 0.28, p = 0.054). Between-group comparison: p = 0.0011. In the chronic stage (days 21–29), Bi-HPC events were longer (17.81 s) than isolated ones (5.93 s; Mann–Whitney p < 0.0001, r = 0.38). (M) CH amplitude: both event types increased (isolated +8.68 µV·day⁻¹, R² = 0.50, p = 0.0050; Bi-HPC +12.37 µV·day⁻¹, R² = 0.54, p = 0.0029); no significant difference between groups (p = 0.38). (N) EF duration: regionally isolated events increased moderately (+0.115 s·day⁻¹, R² = 0.57, p = 0.0017), while Bi-HPC events increased steeply (+0.646 s·day⁻¹, R² = 0.73, p = 0.0001). Between-group p < 0.0001; in the chronic stage, Bi-HPC events were longer (24.82 s vs. 11.46 s; p < 0.0001, r = 0.33). (O) EF amplitude: both event types rose sharply (isolated +7.93 µV·day⁻¹, R² = 0.93; Bi-HPC +9.12 µV·day⁻¹, R² = 0.82; both p < 0.0001). Between-group p = 0.39; in the chronic stage, Bi-HPC events showed slightly lower amplitudes (324 µV vs. 335 µV; p = 0.0061, r = 0.05). P-Q | Early hemispheric asymmetry predicts later seizure burden. Bar plots show LOSO AUC for models trained on week1 (days < 7) metrics to classify high vs. low week4 (days > 21) seizure rate. Dashed line = chance (AUC = 0.5). Numbers above bars indicate permutation p-values (one-sided, AUC > 0.5; 5,000 shuffles) with FDR-corrected q-values. Colors group features (Spikes / FRs / Centrality / Lateralization). IHK mice, n = 13. (P) CH: top feature = global spike lateralization index (LI spikes; AUC ≈ 0.74; p = 0.092; q = 0.184; trend). Other metrics performed near chance (p ≥ 0.62, q ≥ 0.73). (Q) EF: top feature = global fast-ripple lateralization (LI FRs; AUC ≈ 0.85; p = 0.037; q = 0.075; trend after FDR). Other spike or FR features were nonsignificant (p ≥ 0.20, q ≥ 0.41).

However, longitudinal characterization of iPDs revealed a striking hemispheric divergence. In CH, iPDs evolved slowly, with modest but significant increases in duration and amplitude across days, whereas rate showed only an uprise trend that did not reach statistical significance (Fig. 4D–E, H–I). The mean ISI within each event did not significantly change with time. In contrast, EF displayed a clear, cumulative amplification of all temporal and morphological features over time - iPDs became more frequent, longer, contained more spikes per event, and those spikes were larger in amplitude (Fig. 4F–G, J–K). Thus, as expected, pathological discharge dynamics strengthen preferentially within the kainate injected hemisphere.

We then compared unilateral versus bilateral hippocampal paroxysmal discharges, corresponding to iPDs in both hippocampi with an overlap of at least 1 spike. Across both hemispheres, bilateral iPD tended to be longer than iPD occurring only in one hippocampus already in the early latent phase. The duration of bilateral iPD then exhibited a steep temporal escalation across time, while isolated events remained short (Fig. 4L–O).

Next, we asked whether early interictal activity predicts chronic seizure burden. Week1 spike rates showed no detectable association with week4 seizure frequency in either hemisphere (Fig. 4B–C), suggesting that early spiking alone does not determine later ictogenesis.

Finally, we tested whether early electrophysiological asymmetries could predict later seizure severity using a machine-learning analyses based on LOSO scoring of epileptiform features (see M&M). LOSO analysis showed that week1 lateralization indices - especially fast-ripple and spike dominance (for CH and EF respectively) - were the strongest predictors of high week4 seizure rates, outperforming absolute spike or HFO rates alone (Fig. 4P–Q). Thus, early hemispheric imbalance, rather than overall activity load, may serve as a biomarker of future seizure burden.

Together, these results reveal a biphasic trajectory: seizures emerge bilaterally, but subsequent pathological activity consolidates within the EF, where paroxysmal discharges progressively intensify and early HFO/spike lateralization carries prognostic value.

### Rapid large-scale co-spiking emergence

To determine how DG-related epileptic activity engages distributed brain regions during epileptogenesis, we quantified short-latency co-spiking between EF spikes and distributed cortical and hippocampal regions. mSEEG recordings revealed synchronous spike discharges across hippocampal and neocortical areas as illustrated in the example taken from a d5 recording in Fig. 5A, indicating rapid large-scale recruitment (Fig. 5A). Anchor-based co-spiking analysis showed that, by Week 1 (W1), EF spikes already triggered significant short-latency responses in multiple contralateral prefrontal, motor, and visual regions, as well as ipsilateral subiculum and medial entorhinal cortex (Fig. 5B). Thus, widespread long-range co-spiking is established early after injury. Whereas DG-driven co-spiking was widespread and bilateral at Week 1, by Week 4 it persisted in a more selective manner, involving fewer and predominantly ipsilateral cortical regions, notably prefrontal, secondary motor, and visual areas.

**Figure 5:**
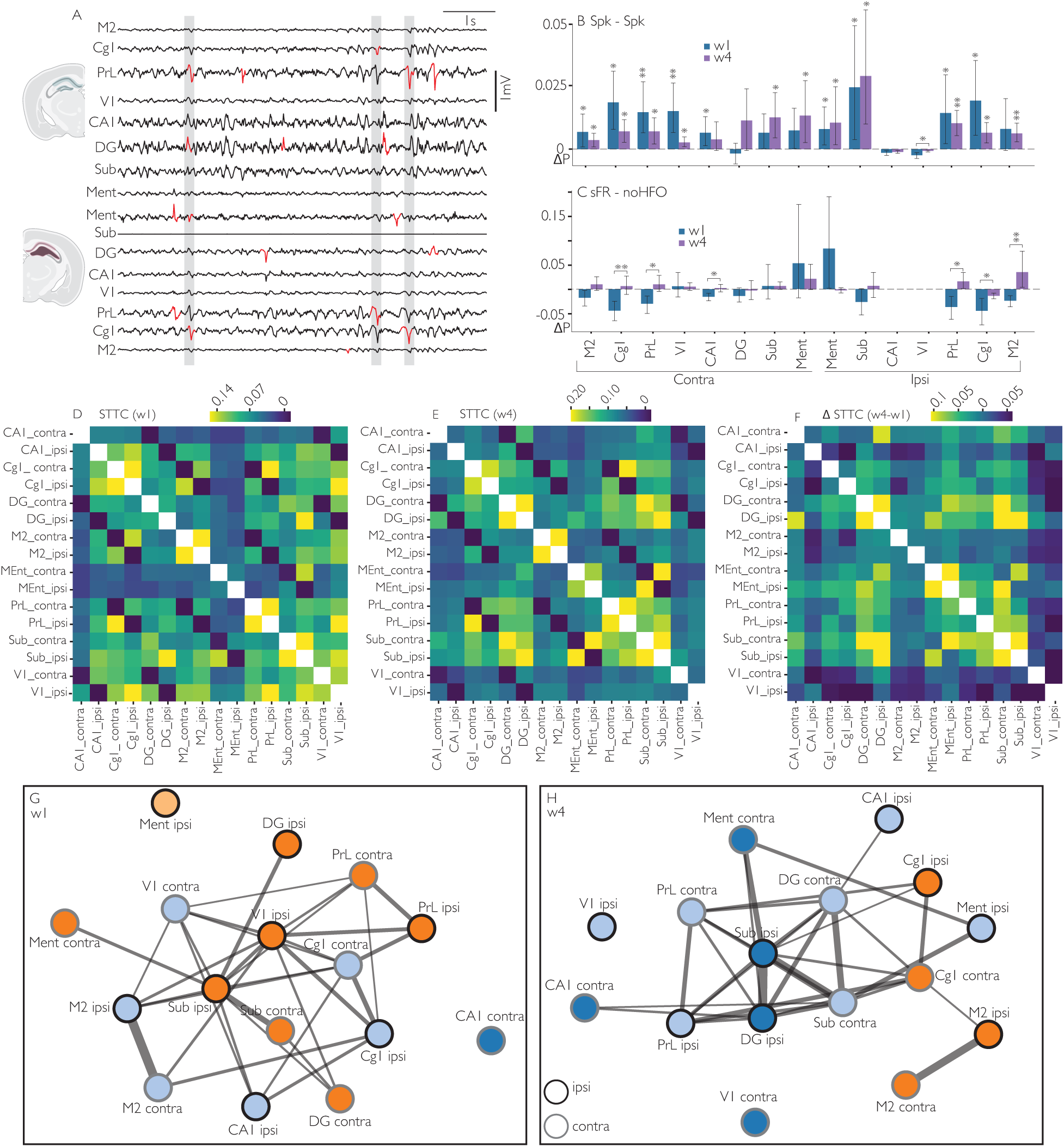
Short-latency dentate gyrus co-spiking rapidly engages cortical networks and becomes more integrated over time. A | Representative EEG traces illustrating synchronous epileptic discharges across distributed cortical and hippocampal regions at d5. Red markers denote detected spikes; grey shaded windows (80 ms) indicate the coincidence window used to define co-occurrence. Spikes occurring within the same window across different regions were classified as synchronous. B | Anchor-based co-spiking probability (ΔP, 0–20 ms). Bars show mouse-level mean ΔP (probability that a spike in the anchor region EF is accompanied by ≥1 spike in the target region within 0–20 ms, minus a ±50 ms jitter baseline). Error bars are bootstrap 95% confidence intervals across mice. One-sided permutation tests against zero (H₁: ΔP > 0) were performed per region and week, with Benjamini–Hochberg FDR correction across regions. Week 1 (W1): significant ΔP > 0 in CA1_contra (p_FDR = 0.028), Cg1_contra (p_FDR = 0.011), Cg1_ipsi (p_FDR = 0.018), M2_contra (p_FDR = 0.048), Ment_ipsi (p_FDR = 0.030), PrL_contra (p_FDR = 0.006), PrL_ipsi (p_FDR = 0.034), Sub_ipsi (p_FDR = 0.042), and V1_contra (p_FDR = 0.006). Week 4 (W4): significant ΔP > 0 in Sub_ipsi (p_FDR = 0.0060), PrL_ipsi (p_FDR = 0.017), M2_ipsi (p_FDR = 0.019), Cg1_ipsi (p_FDR = 0.021), and V1_ipsi (p_FDR = 0.021). Paired permutation tests (H₁: ΔP_W4 > ΔP_W1) revealed a significant increase only in V1_ipsi (p_FDR = 0.0375). Summary: Co-spiking with EF was already widespread at W1, involving contralateral prefrontal (PrL, Cg1), motor (M2), and visual (V1) cortices, and by W4 persisted primarily in ipsilateral subiculum and visual cortex. The only significant temporal increase occurred in V1_ipsi, suggesting progressive recruitment of ipsilateral sensory areas into the epileptogenic co-spiking network. C | FR-conditioned co-spiking contrast (ΔP_FR–none, 0–20 ms). Bars show mouse-level mean difference in co-spiking probability between fast ripple (FR)-tagged anchor spikes and non-FR anchor spikes (none). Positive values indicate that FR-tagged spikes in EF were more effective in eliciting co-spikes in the target region within 0–20 ms than other spikes. Error bars show bootstrap 95% CIs across mice. No region showed significant ΔP_FR–none > 0 after FDR correction in either week. Paired permutation tests (H₁: ΔP_FR–none_W4 > ΔP_FR–none_W1) revealed significant increases in M2_ipsi (p_FDR = 0.0030), Cg1_contra (p_FDR = 0.0067), PrL_contra (p_FDR = 0.0115), CA1_contra (p_FDR = 0.0135), and Cg1_ipsi (p_FDR = 0.0470). Summary: At W1, FR-tagged spikes were slightly less co-spike-effective than non-HFO spikes, but by W4 this difference was attenuated or reversed, indicating a relative normalization of co-spiking efficiency and emergence of network-level autonomy, with large-scale synchronization no longer dependent on focal HFOs. D–F | Spike-time tiling coefficient (STTC) connectivity matrices (τ = 35 ms). Heatmaps show mean STTC across mice for W1 (D) and W4 (E), and their difference (F, W4–W1). The ΔSTTC map highlights regions where short-latency co-variations strengthened over time. STTC was computed pairwise for all regions (diagonal = 0); color scales span the 5th–95th percentiles of observed values. G–H | Graph-level organization of short-latency synchrony at W1 and W4. Nodes represent cortical and hippocampal regions; edge width is proportional to the mean STTC strength between connected pairs; Node fill color denotes Louvain community (module), whereas node outline denotes hemisphere (black = ipsilateral, grey: contralateral). At W1 (G), highly modular organization dominated by bilateral hippocampal–prefrontal coupling was observed, with dense intra-hemispheric links. At W4 (H), network density increased, accompanied by a reorganization of community structure and strengthened interhemispheric and sensorimotor interactions.

To test whether high-frequency oscillations facilitate propagation, we compared co-spiking following fast ripple–tagged versus non-HFO DG spikes. No region exhibited significantly greater co-spiking for FR-tagged spikes at either timepoint (Fig. 5C), indicating that large-scale synchrony does not rely on HFO-associated events. However, several cortical and hippocampal regions showed a significant reduction in the difference between FR- and non-HFO–conditioned co-spiking from W1 to W4, suggesting that as the network matures, synchronization becomes increasingly independent of specific focal spike features and more autonomously coordinated.

Pairwise Spike-Time Tiling Coefficient (STTC) analysis revealed a convergent temporal reorganization of network synchrony. At W1, synchrony was structured into partially segregated hippocampal–prefrontal modules (Fig. 5D, G). By W4, overall synchrony increased, interhemispheric homotypic interactions strengthened, resulting in a denser and more integrated large-scale network, with the strongest interactions between the DG and subiculum regions (Fig. 5E–H). Together, these findings demonstrate that short-latency EF-driven co-spiking emerges rapidly, is initially widespread and bilaterally distributed, and reorganizes into a more cohesive cortical–hippocampal synchrony network, with increasing involvement of the ipsilateral visual cortex during epileptogenesis.

### Symmetric EF–CH coupling during IEDs

Cross-correlation analyses of interictal spikes (IS + ST) further illustrate that, already in the early latent stage (W1), most epileptic spikes tend to co-occur in close proximity between the 2 hippocampi, revealing a near-synchronous activation between the EF and CH (Fig. 6A–B). The population cross-correlograms were centered around zero lag with no systematic bias, indicating a high degree of temporal symmetry between the two hemispheres. Although CH spikes tended to occur slightly before EF (median lag_COM ≈ −2 ms), this small offset was not statistically significant.

**Figure 6:**
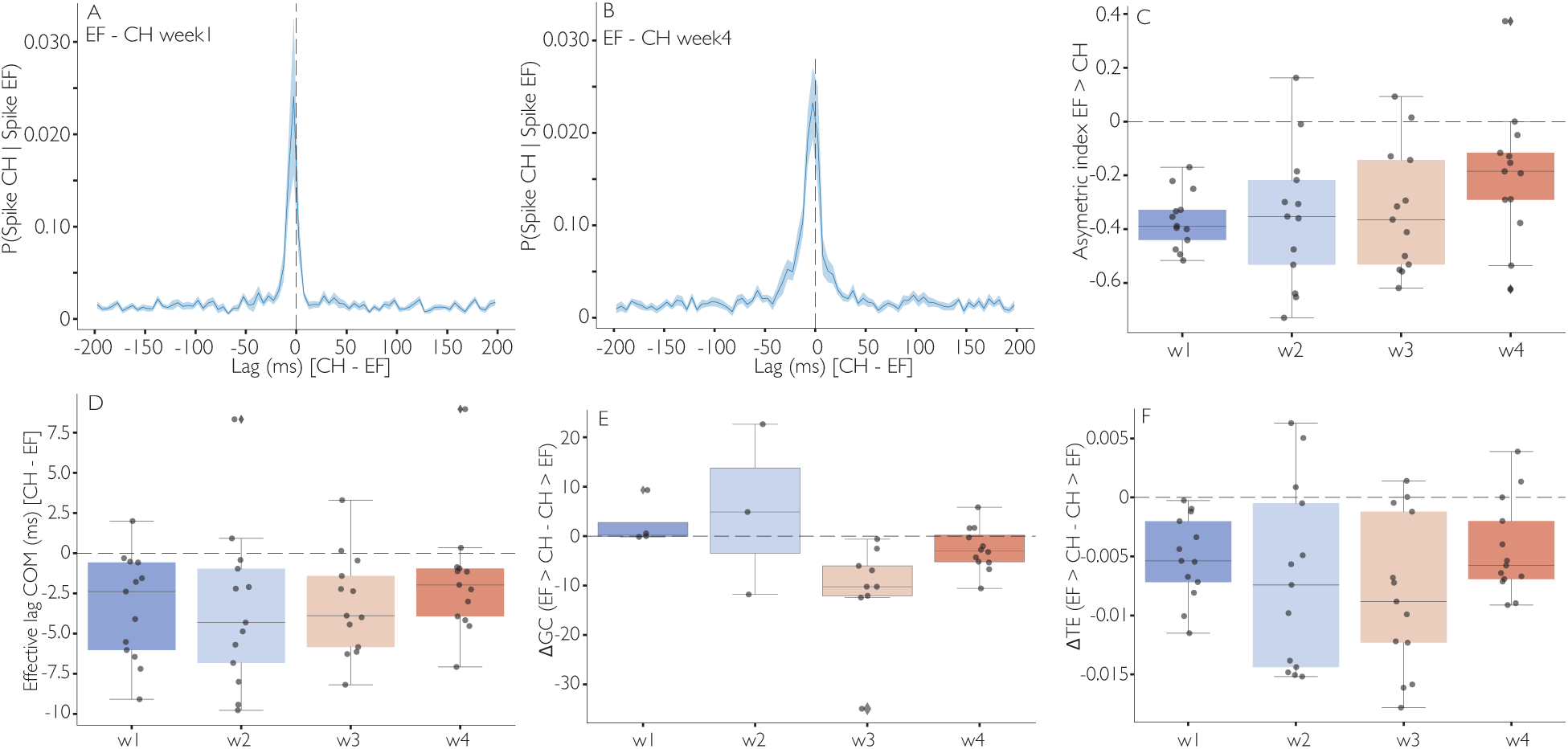
Directional dynamics and causal coupling between the epileptogenic and contralateral dentate gyrus during epileptogenesis. A–B | Spike cross-correlograms between EF and CH (reference = EF, target = CH) averaged across mice at week 1 (A) and week 4 (B). The y-axis indicates the probability of observing a CH spike in each lag bin (5 ms) conditional on a EF spike at t = 0. Shaded areas represent SEM across animals. The central peak remained centered around 0 ms throughout the latent period, indicating near-synchronous activation of both dentate gyri. C | Asymmetry index quantifying the relative strength of short-latency correlations in the positive (0–50 ms) versus negative (−50–0 ms) lag windows. Negative values indicate that CH spikes tend to precede EF spikes. The asymmetry index was significantly negative during the first week (median = −0.39, p = 0.0002 vs 0, Wilcoxon test) and gradually increased toward zero across weeks (linear regression: slope = +0.05·week^-1^, p = 0.055). A paired comparison confirmed a significant increase between week 1 and week 4 (p = 0.02, Wilcoxon signed-rank). D | Effective lag (center-of-mass of the cross-correlation within ±50 ms), indicating the average temporal offset between CH and EF spikes. The lag remained close to zero across weeks (median ≈ −2 ms; linear regression: slope = +0.53 ms·week^-1^, p = 0.27; paired Wilcoxon W1 vs W4, p = 0.41), consistent with quasi-synchronous coactivation. E | Directional Granger causality (ΔGC = GC_EF→CH − GC_CH→EF) computed on binned interictal spike trains (10–20 ms bins, 1 s windows) showed no consistent directional bias (linear regression: slope = −1.21 GC·week^-1^, p = 0.25). F | Directional transfer entropy (ΔTE = TE_EF→CH − TE_CH→EF) computed on the same spike trains also revealed no significant trend (linear regression: slope = −0.001 TE·week^-1^, p = 0.91; paired Wilcoxon W1 vs W4, p = 0.38). Boxplots represent median ± interquartile range; dots show individual mice. Together, these results indicate that the EF–CH interaction remains dominated by fast, reciprocal coupling that becomes progressively more symmetric during epileptogenesis.

The asymmetry index (reflecting the relative temporal precedence of spikes between EF and CH) was significantly negative at week 1 and progressively approached zero across weeks (Fig. 6C), reflecting a gradual equalization of short-latency interactions between both DG rather than a unidirectional propagation from the epileptogenic side. The effective lag (defined as the center-of-mass of the cross-correlogram) remained close to zero throughout the latent phase (Fig. 6D). Granger causality and transfer entropy analyses on binned interictal spike (spike counts per fixed temporal window) (Fig. 6E–F) showed no consistent directional bias across weeks (Supplementary Results), confirming that the EF–CH coupling remained bidirectional and largely symmetric during epileptogenesis.

### Generalized spikes mark network consolidation

Building on these observations of progressively organized inter-regional synchronization, we next turned our attention to generalized interictal spikes (GS), defined as synchronous spike events detected in at least three distinct recording bundles within an 80 ms temporal window, to assess how their dynamics recapitulate the consolidation of the epileptic network. While already present during the early latent phase, GS became a prominent, increasingly frequent feature of the recordings by the chronic stage. Representative waveforms illustrate their spatially widespread and synchronous nature, engaging both hippocampal and neocortical sites (Fig. 7A). Quantification of GS occurrence revealed an increase across nearly all recorded regions (Fig. 7B). While early GS were largely restricted to hippocampal structures, by the third and fourth weeks they had become frequent in associative cortices such as Cg1, PrL, and M2, consistent with the large-scale recruitment of cortical nodes into the epileptic network. We then analyzed the spectral components of GS by assessing their co-occurrence with HFOs. The proportion of GS associated with FRs rose markedly over time in both hippocampal and cortical regions, whereas GS associated with RPs also increased but showed a broader distribution across the network (Fig. 7C). This pattern suggests a progressive embedding of pathological HFOs within generalized events, particularly FRs, which are strongly linked to epileptogenic processes.

**Figure 7:**
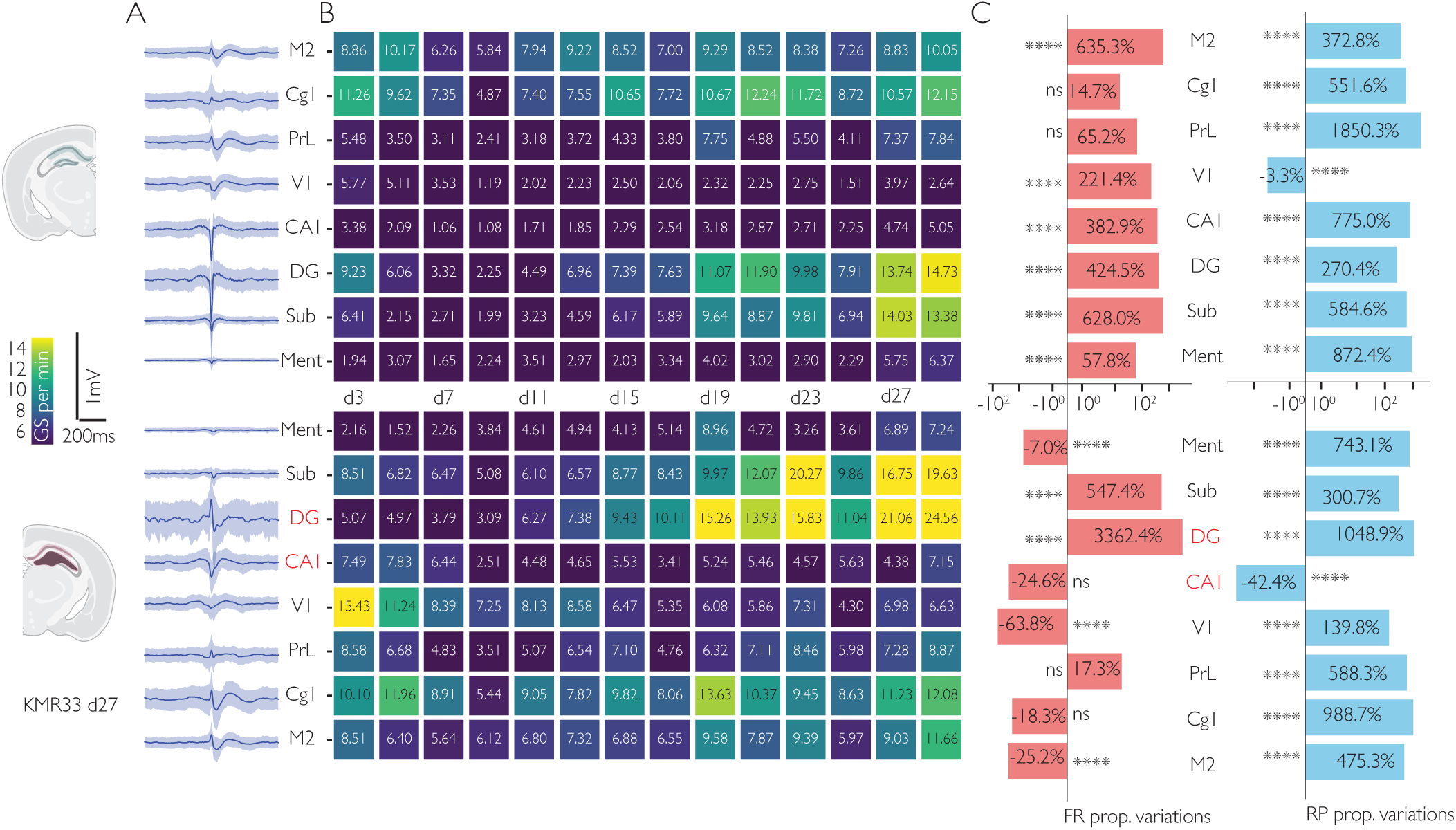
Generalized interictal spikes (GS) mirror the progressive reorganization of the epileptic network. A | Average GS waveforms recorded during the chronic phase in a representative epileptic mouse. Each trace corresponds to one recorded region (M2, Cg1, PrL, V1, CA1, DG, Sub, Ment), with shaded envelopes representing the standard deviation across events. The spatially widespread and synchronous character of GS is evident across cortical and hippocampal regions. B | Heatmaps showing mean GS occurrence rates (events·min^-1^) across the latent phase (days 3–27 post–SE). Each cell represents the group average for one region at a given time point, color-coded from low (purple) to high (yellow/green) GS activity. Early after SE, GS were infrequent and regionally restricted but progressively increased in both hippocampal and neocortical sites over time. C | Bar plots quantifying relative changes in GS-associated high-frequency oscillations (HFOs) between Week 1 and Week 4. Red bars indicate variations in the proportion of GS coupled to fast ripples (sFR), and blue bars indicate variations in GS coupled to ripples (sRP). Percent changes are displayed for each recorded region, with significance levels (**** p < 0.0001, etc.) denoting paired comparisons. While sFR generally increased in hippocampal and cortical nodes, ripple-associated GS tended to expand more broadly across the network. Together, these analyses show that GS evolve from relatively weak, regionally confined events to robust, bilaterally distributed discharges associated with HFOs, thereby recapitulating the large-scale reorganization of the epileptic network during the latent and chronic phases.

Together, these findings show that GS evolve from sporadic discharges confined to a small set of regions to highly synchronized events spanning multiple cortical and hippocampal regions. Their association with pathological HFOs further underscores GS as a hallmark of large-scale network consolidation during the latent phase, foreshadowing the emergence of chronic epilepsy.

### Early EF silencing does not halt progression of spiking activities in the bi-hippocampal network

We next examined whether focal reduction of excitation in the EF during the early latent phase (d2-d7) could mitigate the subsequent progressive increase in spiking activity by using an inhibitory DREADD construct expressed in glutamatergic neurons and CNO administered in the drinking water. KA-injected animals in which the EF modulation treatment was applied (DFIHK group, n=9) showed a robust rise in spike rate and amplitude within the EF across the latent phase, similar to what was observed in epileptic animals without silencing (IHK group, n=12; same group as in previous sections.) (Fig. 8A–B). Regression slopes were comparable between groups, indicating that local silencing did not halt the overall escalation of epileptic activity.

**Figure 8:**
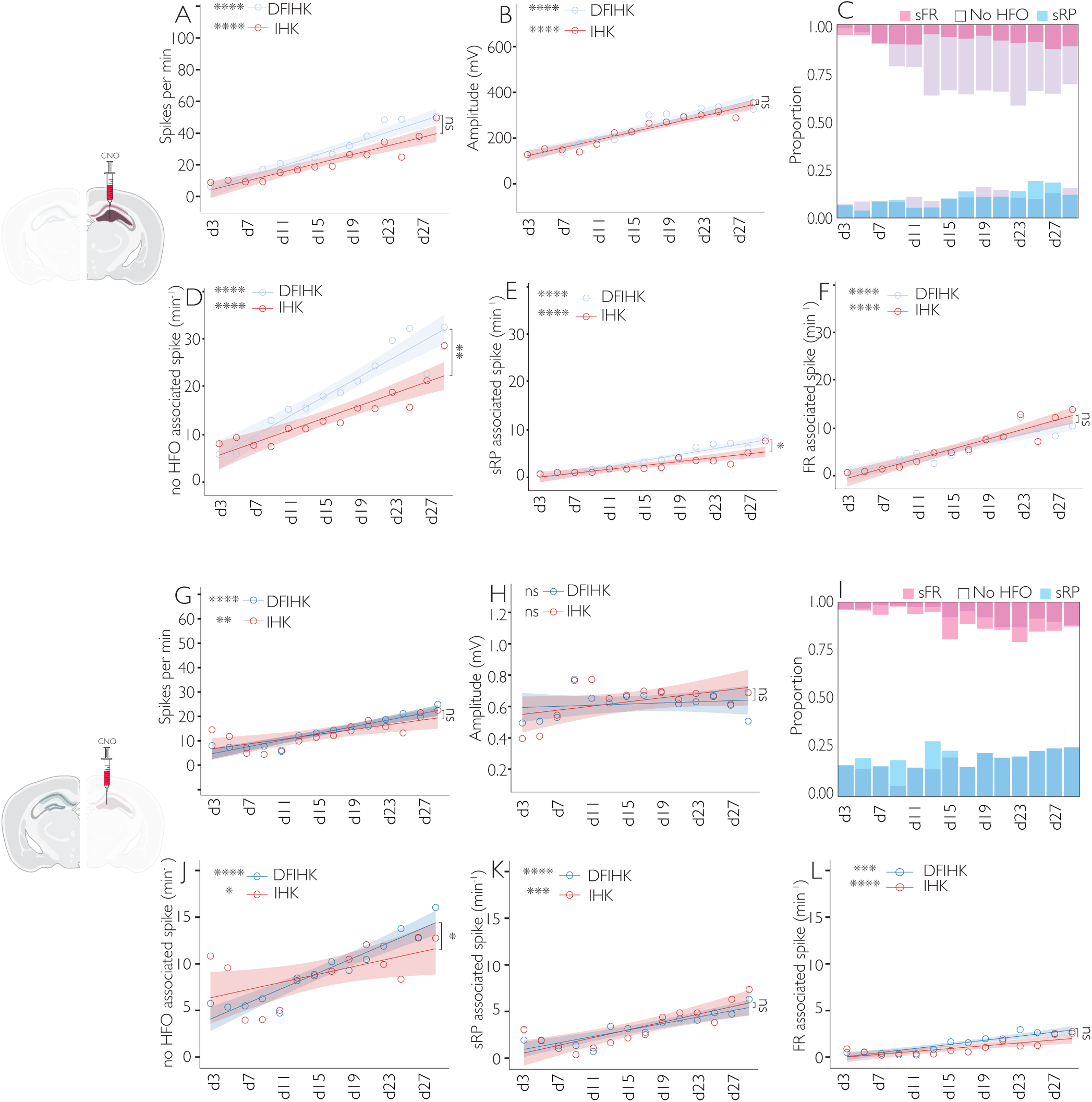
Focal silencing during the latent phase does not alter the global progression of epileptic spiking activity. A | Linear regression of spikes per minute in the EF for epileptic (IHK) animals (red; slope = 1.3640, R² = 0.8655, p < 0.0001) vs. focally silenced animals (DFIHK) (blue; slope = 1.7633, R² = 0.9195, p < 0.0001). B | Linear regression of spike amplitude in the EF for IHK animals (red; slope = 8.6305, R² = 0.9363, p < 0.0001) vs. DFIHK animals (blue; slope = 9.3429, R² = 0.9202, p < 0.0001) C | Proportion of each spike type—noHFO, RP, FR—in the EF over time, comparing IHK vs. DFIHK animals. D | Linear regression of EF spikes without HFO (noHFO) per minute in IHK (red; slope = 0.6476, R² = 0.8197, p < 0.0001) vs. DFIHK (blue; slope = 1.0303, R² = 0.9070, p < 0.0001). Difference is significant (p = 0.0031). E | Linear regression of EF sRP per minute in IHK (red; slope = 0.2042, R² = 0.7622, p < 0.0001) vs. DFIHK (blue; slope = 0.3114, R² = 0.9232, p < 0.0001). Difference is significant (p = 0.0105). F | Linear regression of EF sFR per minute in IHK (red; slope = 0.5114, R² = 0.8915, p < 0.0001) vs. DFIHK (blue; slope = 0.4198, R² = 0.8577, p < 0.0001). Difference is not significant (p = 0.1993). G | Linear regression of spikes per minute in the CH for IHK (red; slope = 0.4898, R² = 0.5313, p = 0.0031) vs. DFIHK (blue; slope = 0.6791, R² = 0.8976, p < 0.0001). H | Linear regression of spike amplitude in the CH for IHK (red; slope = 6.5876, R² = 0.2348, p = 0.0791) vs. DFIHK (blue; slope = 1.7875, R² = 0.0337, p = 0.5297). I | Proportion of each spike type—noHFO, sRP, sFR in the CH over time, comparing IHK vs. DFIHK. J | Linear regression of CH spikes without HFO (noHFO) per minute in IHK (red; slope = 0.2026, R² = 0.3330, p = 0.0307) vs. DFIHK (blue; slope = 0.3973, R² = 0.8980, p < 0.0001). Difference is significant (p = 0.0330). K | Linear regression of CH spikes with sRP per minute in IHK (red; slope = 0.2070, R² = 0.6842, p = 0.0003) vs. DFIHK (blue; slope = 0.1724, R² = 0.8072, p < 0.0001). Difference is not significant (p = 0.4641). L | Linear regression of CH spikes with sFR per minute in IHK (red; slope = 0.0759, R² = 0.6388, p = 0.0006) vs. DFIHK (blue; slope = 0.1095, R² = 0.8791, p < 0.0001). Difference is not significant (p = 0.0963).

Analysis of spike subtypes in the EF revealed a similar distribution between groups, with spikes progressively shifting from being predominantly uncoupled from high-frequency oscillations (HFOs) to being increasingly associated RP and, to a lesser extent, FR (Fig. 8C). Quantitative analysis showed that the rate of noHFO-spikes and RP-associated spikes increased more steeply in silenced animals (Fig. 8D–E), whereas FR-associated spikes followed parallel trajectories across groups (Fig. 8F).

In the CH, both groups exhibited a progressive rise in spike rates (Fig. 8G), while spike amplitudes remained relatively stable (Fig. 8H). The distribution of spike subtypes in the CH mirrored the EF, with noHFO and RP-associated spikes increasing steadily and FR-associated spikes remaining less frequent overall (Fig. 8I–L). Notably, silenced animals displayed a modest but significant increase in the rate of noHFO spikes compared to non-silenced controls (Fig. 8J), whereas RP- and FR-associated spikes showed no group differences (Fig. 8K–L).

Taken together, these results indicate that a focal reduction in glutamatergic neuron activity in the EF during the very first days of the latent phase does not prevent the amplification and network-wide spread of epileptic spiking. Instead, silencing may induce subtle shifts in spike subtype dynamics, particularly by enhancing the rate of spikes uncoupled from HFOs, but the global epileptic trajectory remains unaffected.

### CH silencing accelerates epileptogenesis

To determine whether activity in the CH modulates epileptogenesis, we applied the same chemogenetic modulation of glutamatergic neurons in the CH during the early latent phase in another group of mice (DCIHK, n=9). As shown in Fig.9A-B, spike rate in the CH similarly increased over time in treated (DCIHK) and non-treated animals (IHK), while spike amplitudes remained stable and comparable (Fig. 9B). The distribution of spike subtypes—spikes without HFOs, ripple-associated spikes (RP), and fast ripple–associated spikes (FR)—was also similar across groups (Fig. 9C–F), indicating that contralateral silencing did not substantially affect local CH dynamics.

**Figure 9:**
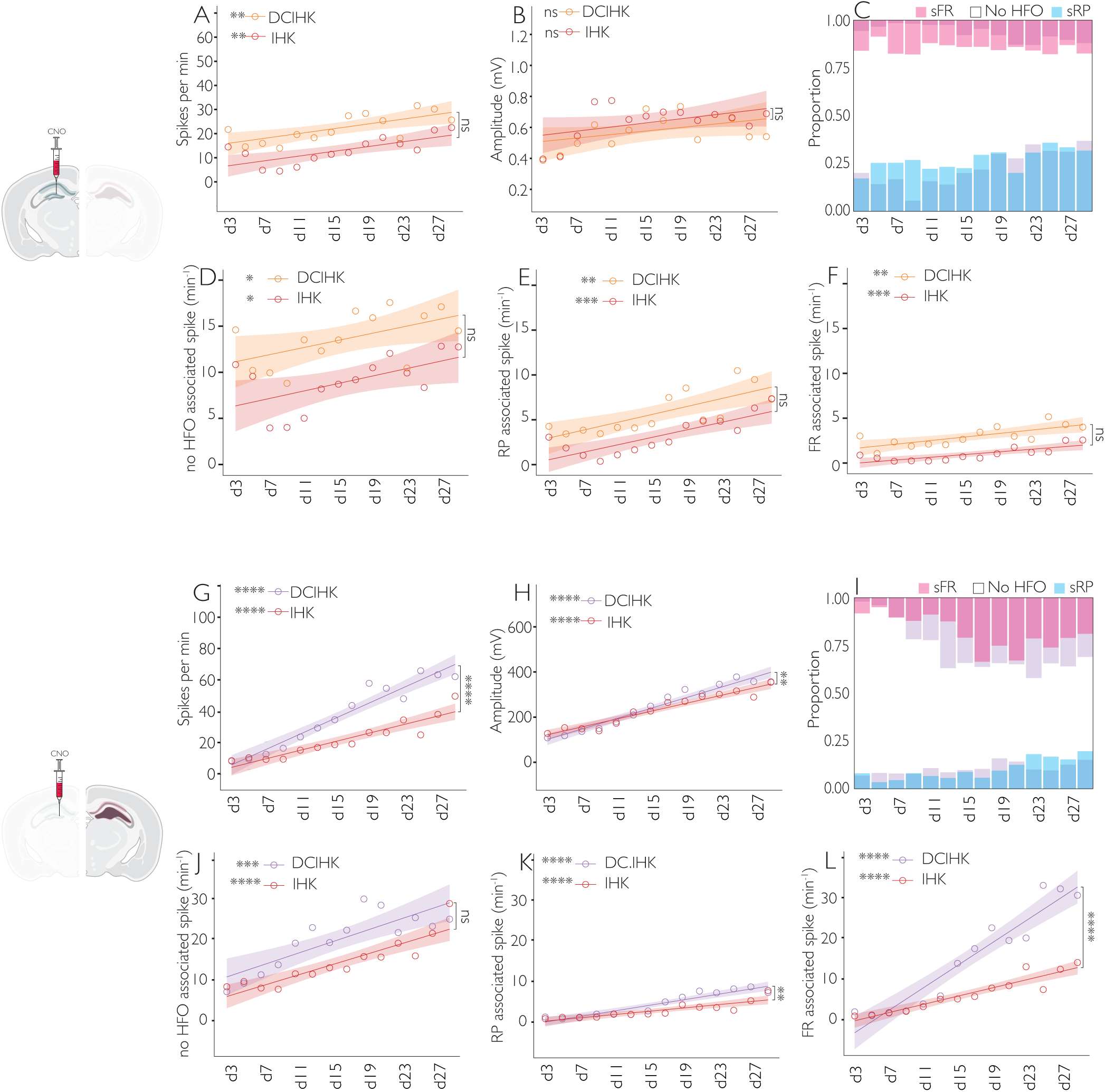
Contralateral silencing during the early latent phase worsens epileptic activity. A | Linear regression of spikes per minute in the CH for epileptic (IHK) animals (red; slope = 0.4898, R² = 0.5313, p = 0.0031) vs. contralaterally silenced (DCIHK) animals (orange; slope = 0.5045, R² = 0.5193, p = 0.0036). B | Linear regression of spike amplitude in the CH for IHK (red; slope = 6.5876, R² = 0.2348, p = 0.0791) vs. DCIHK (orange; slope = 6.3312, R² = 0.2435, p = 0.0730). C | Proportion of different spike types—e.g., noHFO, sRP, sFR - in the CH over time, comparing IHK vs. DCIHK. D | Linear regression of CH spikes without HFO (“noHFO”) per minute in IHK (red; slope = 0.2026, R² = 0.3330, p = 0.0307) vs. DCIHK (orange; slope = 0.1934, R² = 0.3088, p = 0.0391). The difference is not significant (p = 0.9377). E | Linear regression of CH sRP per minute in IHK (red; slope = 0.2070, R² = 0.6842, p = 0.0003) vs. DCIHK (orange; slope = 0.2184, R² = 0.5886, p = 0.0014). The difference is not significant (p = 0.8649). F | Linear regression of CH spikes with sFR per minute in IHK (red; slope = 0.0759, R² = 0.6388, p = 0.0006) vs. DCIHK (orange; slope = 0.0992, R² = 0.5578, p = 0.0021). The difference is not significant (p = 0.4434). G | Linear regression of spikes per minute in the EF for IHK (green; slope = 1.3640, R² = 0.8655, p < 0.0001) vs. DCIHK (orange; slope = 2.4668, R² = 0.9360, p < 0.0001). H | Linear regression of spike amplitude in the EF for IHK (green; slope = 8.6305, R² = 0.9363, p < 0.0001) vs. DCIHK (orange; slope = 11.4252, R² = 0.9509, p < 0.0001). I | Proportion of each spike type—noHFO, sRP, sFR - in the CH over time, comparing IHK vs. DCIHK. J | Linear regression of EF spikes without HFO per minute in IHK (red; slope = 0.6476, R² = 0.8197, p < 0.0001) vs. DCIHK (purple; slope = 0.7135, R² = 0.6924, p = 0.0002). No significant difference (p = 0.6860). K | Linear regression of EF s RP per minute in IHK (red; slope = 0.2042, R² = 0.7622, p < 0.0001) vs. DCIHK (purple; slope = 0.3447, R² = 0.8934, p < 0.0001). Difference is significant (p = 0.0032). L | Linear regression of EF sFR per minute in IHK (red; slope = 0.5114, R² = 0.8915, p < 0.0001) vs. DCIHK (purple; slope = 1.4016, R² = 0.9160, p < 0.0001). Difference is significant (p < 0.0001). All linear regressions display shaded areas indicating the standard error of the mean (SEM).

In contrast, strong effects were observed within the epileptogenic focus (EF). Mice with contralateral silencing displayed markedly higher spike rates and a steeper increase over time compared with KA controls (Fig. 9G). Spike amplitudes were also significantly larger in silenced animals (Fig. 9H). Subtype analysis revealed that while the rate of noHFO spikes rose similarly across groups (Fig. 9J), silenced animals exhibited a clear increase in ripple-associated spikes (Fig. 9K) and a robust enhancement of fast ripple–associated spikes (Fig. 9L).

Together, these results demonstrate that silencing the contralateral hippocampus during the early latent phase does not dampen epileptogenesis but instead amplifies it, leading to higher spiking rates and stronger pathological high-frequency components in the epileptogenic focus. This suggests that the contralateral hippocampus may exert a modulatory or compensatory influence that normally restrains the escalation of epileptic discharges.

## Discussion

### From focal to distributed epileptogenesis

Our data reveal that epileptogenesis in the intrahippocampal KA model is a dynamic process involving a gradual transition from local to distributed network activity. In the early latent phase, epileptiform discharges were restricted to the ipsilateral hippocampus, particularly within the dentate gyrus and CA1, consistent with the initial lesion site and its strong intrinsic connectivity (Lévesque & Avoli, 2013; Sheybani et al., 2018). Over time, the activity propagated to the contralateral hippocampus and then to distant cortical regions, resulting in the establishment of a bilateral epileptogenic network.

This progression supports the view that epileptogenesis reflects a network maturation process rather than a simple spread of pathology. Early alterations within the lesioned hippocampus likely serve as a “seed” that entrains connected areas via recurrent excitatory projections and reduced inhibitory control. Similar transitions have been described in chronic imaging studies showing a shift from localized to widespread activation patterns during seizure onset in both animal models and human patients (Engel et al., 2013; Spencer, 2002).

The gradual bilateralization of spikes highlights the involvement of commissural and cortical relays in network consolidation. At the interictal level, contralateral recruitment became apparent after a short delay of several days, indicating that secondary plastic changes, rather than passive propagation, underlie interhemispheric integration. Nevertheless, once contralateral spikes emerged, they rapidly engaged in short-latency coactivation with the ipsilateral focus, consistent with the early bilateral co-spiking observed in Fig. 5.

### Microcircuit reorganization and HFO biomarkers

At the focal level, the increasing occurrence of high-frequency oscillations (HFOs) associated with epileptiform spikes provides evidence for the maturation of pathological microcircuits. Ripples and fast ripples have been widely recognized as biomarkers of epileptogenicity, reflecting abnormal synchronization among small clusters of pyramidal cells (Bragin et al., 1999; Jelerys et al., 2012).

Our data show that spike-related HFOs emerge in the ipsilateral hippocampus before appearing in cortical regions, suggesting a temporal hierarchy: local microcircuit reorganization precedes large-scale synchronization. These findings are consistent with the notion that epileptogenesis may originate from local microcircuit dysfunctions within the epileptogenic focus, which have been associated in previous studies with processes such as mossy fiber sprouting, altered dendritic integration, or interneuron loss (Nadler, 2003, Bernard et al., 2004, Maglóczky & Freund, 2005). Although these cellular mechanisms were not directly assessed here, they provide a plausible substrate for the progressive large-scale synchronization observed in our recordings.

Furthermore, the increase in spike amplitude and HFO frequency during the latent phase points to a process of activity-dependent potentiation, possibly involving enhanced NMDA receptor function, axonal sprouting, or astrocytic dysregulation (Scharfman, 2007; Vezzani et al., 2011). Together, these findings underscore the interdependence of microcircuit plasticity and network-level coupling during epileptogenesis.

### Contralateral and cortical recruitment reflect network-level propagation

The delayed yet robust emergence of synchronous discharges in the contralateral hippocampus suggests a form of secondary epileptogenesis, wherein distant but interconnected regions become progressively co-opted into the epileptic network. However, the predominance of ripple-frequency HFOs and the exacerbation of ipsilateral spiking and fast ripples upon contralateral silencing indicate that the CH remains only partially epileptogenic and may exert a restraining influence. Anatomically, this could result from potentiation of commissural projections between CA3 and CA1, or through intermediary relays such as the entorhinal cortex and subiculum, both of which are heavily interconnected bilaterally (van Strien et al., 2009).

Interestingly, the progressive increase in spike rate and synchronization within cortical regions such as M2, PrL, and Cg1 indicates that higher-order association cortices actively participate in the expanding network. This cortical recruitment may reflect both top-down modulation—through corticohippocampal feedback loops—and bottom-up entrainment from hippocampal generators. Functional imaging in chronic TLE patients has similarly shown that the prefrontal and cingulate cortices become pathologically coupled to mesial temporal structures as the disease progresses (Bartolomei et al., 2017).

Our results indicate that cortical regions become increasingly and persistently engaged in epileptic synchronization as epileptogenesis progresses. While our data do not directly demonstrate autonomous epileptogenicity of cortical nodes, the progressive strengthening of cortico-hippocampal synchrony suggests that these regions may contribute actively to the maintenance of large-scale epileptic dynamics.

### Directional dynamics and bilateral coupling during epileptogenesis

The analysis of short-latency interactions between EF and CH reveals that both hippocampi engage in highly synchronous discharges at early stages of epileptogenesis, characterized by sharp cross-correlation peaks centered around zero lag. This temporal structure indicates that contralateral recruitment unlikely to rely solely from slow propagation but rather from fast commissural or associational pathways capable of coordinating discharges within a few milliseconds (Spencer et al., 1984; Jefferys C’eand Haas, 1982; Avoli et al., 1996).

Over the latent period, the reduction in cross-correlation asymmetry suggests a gradual rebalancing of directional interactions between both sides. Initially, CH spikes showed a slight tendency to precede EF, which could be compatible with a transient contralateral modulation of excitatory drive (Toyoda et al., 2013; Lévesque et al., 2018), although our GC and TE analyses did not reveal a dominant direction of causal influence. With time, this small bias diminished, consistent with the strengthening of commissural feedback and the establishment of a more symmetric, reciprocal network. The near-zero effective lag and the absence of consistent directional bias in Granger causality or transfer entropy further support the view that epileptogenesis promotes bilateral synchronization through mutual entrainment rather than hierarchical drive (Engel et al., 2013; Bartolomei et al., 2017).

These results align with network-based models of temporal lobe epilepsy, in which initially focal hyperexcitability progressively engages homologous regions via long-range plasticity, leading to the formation of bilateral resonant loops (Bragin et al., 2000; Schevon et al., 2007; Keller et al., 2014). Such dynamic equalization of hippocampal interactions may represent a critical step in the transition from a unilateral focus to a distributed epileptic network, facilitating the emergence of large-scale synchronous events and secondary generalization (Bertram, 2013; Jirsa et al., 2017).

### Network resilience to focal inhibition

Chemogenetic silencing experiments provide critical insight into the causal organization of the epileptic network. In our study, chemogenetic inhibition of the ipsilateral or contralateral hippocampus targeted local hippocampal activity but did not prevent the emergence of global synchronization. This resilience suggests that once multiple nodes become integrated, the epileptic network can compensate for local perturbations by reweighting effective connectivity.

Such compensatory recruitment of alternative pathways is consistent with prior reports showing that focal inhibition or surgical resection often fails to abolish seizures in chronic TLE (Krook-Magnuson et al., 2013; Rosenow & Lüders, 2001). From a network perspective, our results are consistent with the idea that epileptogenesis leads to the emergence of a distributed and robust network state, in which partial suppression of individual nodes is insufficient to disrupt global synchronization.

Mechanistically, several processes may underlie this resilience: (1) redundant excitatory projections between hippocampal and cortical areas; (2) homeostatic synaptic scaling maintaining network excitability; and (3) astrocytic and inflammatory feedback promoting persistent synchronization. This plasticity, while maladaptive, likely evolved to preserve information flow under injury, inadvertently stabilizing the epileptic state.

### Mechanistic implications and conceptual model

Taken together, these findings support a model in which epileptogenesis progresses through distinct but overlapping stages: (1) Local initiation, characterized by HFO emergence and increased excitability within the KA focus. (2) Network coupling, where interregional synchrony develops via commissural and cortical pathways. (3) Distributed stabilization, marked by bilateral integration and resistance to focal interventions.

This framework integrates cellular, synaptic, and network mechanisms into a unified view of temporal lobe epileptogenesis. It implies that interventions limited to the primary focus may be insufficient once large-scale coupling has occurred. Future therapeutic strategies should therefore aim to modulate the network as a whole, either through targeted neuromodulation (optogenetic, chemogenetic, or electrical) or by influencing plasticity mechanisms that sustain pathological synchrony.

Finally, our approach—combining chronic multi-site electrophysiology and causal manipulations—highlights the importance of network-level monitoring during the latent phase, when epileptogenesis remains potentially reversible. Understanding how pathological synchrony consolidates across regions may ultimately guide early interventions capable of preventing chronic epilepsy rather than merely suppressing seizures.

### Implications for therapy and early intervention

Our use of chronic multi-site recordings provided a unique mesoscale view of epileptogenesis, bridging the gap between microcircuit studies and whole-brain imaging. However, certain limitations should be noted. First, chemogenetic inhibition was applied during a limited temporal window; longer or repeated modulation could yield different outcomes. Second, while DREADD activation provides spatial precision, it lacks the temporal resolution of optogenetic or electrical interventions, which might uncover transient causal interactions.

Future studies could combine closed-loop detection with region-specific inhibition to dissect the directionality of epileptogenic communication. Moreover, integrating electrophysiology with calcium or voltage imaging could help reveal how local circuit changes manifest at the network level.

### Concluding remarks

Epileptogenesis rapidly reorganizes into a distributed, bilateral network that can compensate for focal suppression, indicating that therapies targeting a single hippocampal node are unlikely to prevent disease progression. Effective intervention in TLE will require network-level strategies—whether diagnostic, pharmacological, or neuromodulatory—implemented early in the latent phase, before large-scale synchronization becomes self-sustaining.

## Supporting information

Supp_Methods_Results

