## Supplementary material for "Progressive bilateral recruitment and resilient network reorganization during temporal lobe epileptogenesis": Supp_Methods_Results

**Supplementary Methods**

Electrode Fabrication

Recordings were obtained from 16 stainless-steel electrodes (PFA-coated, 110 μm diameter; Science Products), arranged into 6 polytrodes. Each polytrode bundle was threaded through a cannula, and individual electrode tips were trimmed to the desired length using a micro ruler (Electron Microscopy Sciences). Electrodes were attached to an Electronic Interface Board (EIB; Neuralynx). Tip impedance was standardized to approximately 400 kΩ by gentle gold plating in a diluted gold solution (Neuralynx), which improves conductivity and ensures recording uniformity.

Kainate Injection

A unilateral TLE mouse model was induced using KA injection (Arabadzisz et al., 2005; Riban et al., 2002). As previously described (Sheybani et al, 2018), mice were anesthetized with isoflurane (3% in $O_2$ at 1 L·min$^{-1}$) and placed in a stereotaxic apparatus. The scalp was shaved, disinfected with povidone-iodine (Betadine), and a small craniotomy (<300 μm in diameter) was made above the left dorsal hippocampus (coordinates relative to Bregma: AP −1.8 mm, ML −1.6 mm, DV 1.9 mm). A total volume of 70 nL kainic acid solution (0.35 nmol, 5 mM in 0.9% NaCl, Tocris Bioscience) was then injected into the targeted region using a Nanoinject 3 device (Drummond) connected to a pulled glass capillary (~15 μm tip diameter). The injection rate was maintained at 10 nL·s$^{-1}$. After injection, the capillary remained in place for 3–4 minutes to allow diffusion. The craniotomy was then cleaned, and the scalp incision sutured. Control animals underwent the same procedure but received 0.9% NaCl instead of KA. Any subsequent surgical procedures, including electrode implantations, were performed at least 24 hours after the KA injection (d0; see Figure 1A).

Multi-region implantation

At d1 post-KA injection, mice were anesthetized using a combination of medetomidine (Dormitor®, 0.5 mg·kg$^{-1}$), midazolam (Dormicum®, 5 mg·kg$^{-1}$), and fentanyl (0.05 mg·kg$^{-1}$). Animals were placed in a stereotaxic frame and maintained at a stable body temperature via a closed-loop heating system. The scalp sutures from the KA injection were removed, and eight bilateral brain regions were targeted for chronic electrode implantation (Fig.1B) based on coordinates from the Paxinos Atlas: secondary motor cortex (M2, AP +1.7 mm, ML ±0.5 mm, DV 0.5 mm), cingulate cortex, area 1 (Cg1, AP +1.7 mm, ML ±0.5 mm, DV 1.3 mm), prelimbic cortex (PrL, AP +1.7 mm, ML ±0.5 mm, DV 1.7 mm), primary visual cortex (V1, AP −2.7 mm, ML ±2.0 mm, DV 0.5 mm), field CA1 of hippocampus (CA1, AP −2.7 mm, ML ±2.0 mm, DV 1.3 mm), dentate gyrus (DG, AP −2.7 mm, ML ±2.0 mm, DV 1.7 mm), Subiculum (S, AP −4.3 mm, ML ±2.8 mm, DV 2.7 mm), entorhinal cortex (Ment,: AP −4.3 mm, ML ±2.8 mm, DV 3.6 mm). Polytrodes were affixed to the skull with a UV-curable dental resin (Ivoclar Vivadent). Two stainless-steel screws (P Technologies) were placed above the cerebellum to serve as reference and ground and connected to the EIB by tungsten wire (Science Products). The electrode assembly and EIB were then secured with dental cement (Paladur). An antidote mixture containing flumazenil (Anexate®, 0.5 mg·kg$^{-1}$), atipamezole (Alzane®, 2.5 mg·kg$^{-1}$), and naloxone (1.2 mg·kg$^{-1}$) was administered to reverse anesthesia. Postoperatively, mice were monitored closely, provided with analgesics (Carprofen, 0.5 mg·kg$^{-1}$ in 300 µL 5% glucose solution), antibiotics (Nopil), and a high-nutrition gel (SAFE) in their cage.

Freely Moving Recordings

Beginning on d3 post-KA injection, neural activity was recorded every other day (for ~1 h per session) until d29 using a Digital Lynx SX acquisition system (Neuralynx) at a 32 kHz sampling rate. Bandpass filtering (0.1–9000 Hz) allowed capture of both local field potentials (LFPs) and multi-unit activity (MUA). Recordings were performed in the afternoon to reduce potential circadian variability. To minimize stress during cable connection, mice received slight isoflurane anesthesia. Recording started 10 min after animals regained normal locomotion, ensuring stable physiological conditions.

Epileptiform Spike Detection

A semi-automated detection of epileptic spikes and epileptic events classification pipeline was created using custom MATLAB (R2022b) scripts, based on a previously published approach (Heining et al., 2019). Recordings were down-sampled to 2 kHz and low-pass filtered at 1 kHz. Channels with visible artifacts or large movement-related noise were manually excluded from the analysis. First, manually marked “background periods” were defined by ~50 artifact-free 2 s segments to calculate baseline mean ± SD (Fig.2A). High-frequency noises were detected and excluded automatically when oscillations exceeding mean ± 5 SD in more than 3 electrode bundle in the 700-1kHz band signal. In the 4-40 Hz filtered denoised signal, events exceeding mean ± 5 SD from the baseline were labeled as spikes. A dead time of 80 ms was imposed before detecting subsequent spikes. To avoid duplicate detection of the same event, spikes identified within 5 ms on different electrodes of the same polytrode were compared, and the smaller peaks were discarded.

High-Frequency Oscillations

High frequency oscillations (HFOs) often accompany epileptiform spikes in both clinical and experimental TLE models (Padmasola et al., 2024; Salami et al., 2014; Sheybani et al., 2018). Because spikes that coincide with an HFO are considered as more specific markers of epileptogenicity than spikes without HFOs (Salami et al., 2014; Wang et al., 2013), we restricted our HFOs analysis to Spikes+ripples (sRP, 80–250 Hz) and Spike+fast-ripples (sFR, 250–500 Hz)—occurring within an 80 ms window (±40 ms) around the peak of each detected spike. HFOs events were identified when the amplitude in the corresponding filtered signal exceeded 2× (SD + baseline mean) for ≥2 consecutive cycles (ripples) or ≥4 consecutive cycles (fast-ripples). The duration was defined from the first to the last oscillation above threshold, and the intrinsic frequency was determined by Fourier transform within their respective frequency band.

Classification of Epileptiform Discharges

Epileptiform discharges were categorized by spiking frequency and event duration (Fig. 1C): Isolated Spikes (IS): Discharges that are not followed or preceded by another spike for ≥1 s. Events containing more than two spikes, each separated by less than 1 second, were considered spike bursts and further classified according to their maximal spike rate into Spike Trains (STs) or Paroxysmal Discharges (PDs). STs were defined as events that never reached a spike rate of 5 spikes·s$^{-1}$. PDs were defined as events that reached a spike rate of ≥5 spikes·s$^{-1}$ for more than 1 second. For each event, the first spike not followed by another spike for at least 1 second marked its termination. Finally, all PDs were visually reviewed by two independent reviewers (MH, CQ), blinded to the experimental group (saline vs. IHK) and post-injection day, for seizure detection. Spikes that occur in more than 5 bundles in a 100 ms time-window are classified as network IED (Padmasola et al., 2024b) and were excluded from regionally isolated spikes analysis. Additionally, an epileptic event in one hippocampus is considered to coincide with a contralateral event if the two events partially or completely overlap in time.

Histology

At the end of the experimental protocol (day 29 post–KA injection), mice were deeply anesthetized with pentobarbital (150 mg·kg$^{-1}$, IP) and perfused transcardially with 0.9% NaCl, immediately followed by 4% paraformaldehyde (PFA) in phosphate-buffered saline (PBS, pH 7.4). Brains were extracted and post-fixed overnight at 4 °C in 4% PFA, then rinsed in PBS and stored in PBS until sectioning. Coronal sections (80 µm thickness) were cut using a vibratome (Leica VT1000S) and collected in PBS. Nuclear counterstaining was performed using DAPI (Drop-n-Stain Everbite with DAPI - Biotium, incubation for 15 minutes at room temperature), followed by several washes in PBS. To assess the expression of DREADD constructs, native mCherry fluorescence was visualized by epifluorescence microscopy (Nikon Eclipse 90i microscope) with appropriate excitation/emission filters. Images were acquired under identical exposure parameters across animals to allow qualitative comparison of expression patterns.

Spike rate heatmap:

For each recording, spikes were grouped into their corresponding events (Fig1C). For each event, the following quantities were computed: event start and end sample, number of spikes, and occurrence of specific high-frequency oscillation (HFO) subtypes (fast ripples, ripples, or no-HFO). Event durations were then expressed in seconds based on the recording sampling rate (2 kHz). Analyses were restricted to the epileptic condition and to events labeled as iPD. For each mouse, day, and electrode, the total number of spikes was divided by the corresponding recording duration (in minutes) to obtain the spike rate (spikes per minute). Spike rates were averaged across mice for each combination of day and electrode. The resulting electrode × day matrix was used to construct a heatmap representation of the average iPD spike rate dynamics over time and across recording sites. The heatmap was generated using seaborn and matplotlib in Python (version 3.11). A “magma” colormap was applied, and the color scale maximum (vmax) was set to the 95th percentile of the data distribution to reduce the influence of outliers. Colorbars represent the average spike rate per minute.

Linear regression Spike rate, Amplitude, Spike related HFO / min:

For each day, condition, and electrode, spike rates, amplitude and spike related HFO rate were aggregated across mice. The group-level mean, standard deviation, and sample size were computed, yielding a dataset of average metrics rates as a function of recording day and electrode. For each electrode and condition, the evolution of metrics rate over days was analyzed by ordinary least-squares linear regression (OLS, SciPy, linregress). Regression slopes, intercepts, $R^2$, p-values, and standard errors were extracted. To visualize the fitted trend, predicted values were computed on a uniform grid of days, and 95% confidence intervals around the regression line were estimated using the Student’s t distribution. When multiple experimental conditions were present, regression slopes were compared across conditions using a Z-test for the difference of slopes. The test statistic was defined as: $Z = \frac{\Delta\beta}{\sqrt{SE_1^2+SE_2^2}}$, where $\Delta\beta$is the slope difference and $SE_1, SE_2$are the corresponding standard errors. Two-tailed p-values were reported for each pairwise comparison. For each electrode, metrics (± standard deviation) were plotted over days with regression lines and shaded confidence intervals.

Proportion of spike related HFOs:

For week-level analyses, recording days were mapped to two bins: Week 1 (days 0–7) and Week 4 (days 21–28). Rows outside these ranges were discarded. For each spike, binary flags were created for fast ripples (FR), ripples (RP), NoHFO, and AnyHFO (FR or RP). Data were aggregated per (cond, mouse, week, elec) to obtain, for each grouping, the total number of spikes and the counts of spikes bearing each HFO label. Proportions were then computed as: $prop_L = \frac{n_L}{n_{spikes}},\ L\ \in \{FR, RP, noHFO\}$. Only groupings with non-zero spike counts contributed to proportions. or each electrode (EF and CH) and each metric ($\mathrm{prop}_{\mathrm{FR}}, \mathrm{prop}_{\mathrm{RP}}, \mathrm{prop}_{\mathrm{NoHFO}}, \mathrm{prop}_{\mathrm{AnyHFO}}$), within-mouse changes from Week 1 to Week 4 were tested using the paired Wilcoxon signed-rank test (two-sided). Mice were included only if they had valid proportion estimates at both weeks (pairwise complete cases). Tests were performed (i) overall with all conditions pooled and (ii) stratified by condition (separate tests within each cond). For each test, the script reports sample size (mice), Week 1 and Week 4 means, Wilcoxon statistic, and p-value Bonferonni corrected.

Half-amplitude width analysis:

To estimate the baseline of each electrode, we used the same background marker file as for spike detection. For each electrode, all background windows were concatenated, and the baseline was defined as the median of the pooled samples. For each spike timestamp, the corresponding electrode trace was extracted in a window of ±75 ms around the spike. The spike peak (maximum absolute deviation from the electrode-specific baseline) was then identified within ±150 ms around the provided timestamp. The half-amplitude level was defined as baseline + 0.5 × (peak − baseline). The first and second crossings of this level, before and after the peak, were located using linear interpolation between adjacent samples. The half-amplitude width (HAW) was calculated as the temporal distance between these two crossings and expressed in milliseconds.

Power spectral density:

All available background markers for each recording were used to extract clean 2-s segments from the raw signal. Segments that extended beyond the recording boundaries were discarded. For each mouse, day, and electrode, power spectral analyses were computed on all valid background windows. For each 2-s segment, the power spectral density (PSD) was estimated using Welch's method (pwelch in MATLAB R2022a) with the following parameters: window length: 1 s Hamming window, overlap: 50%, FFT length: 4096 points, frequency resolution: ~0.49 Hz, output units: $V^2 \cdot Hz^{-1}$. The following frequency bands were defined: δ (1–4 Hz), θ (4–8 Hz), β (12–30 Hz), γ (30–80 Hz), and ripple (80–250 Hz). For each band, the absolute power ($V^2$) was computed by integrating the PSD across the band using MATLAB's bandpower function. In addition, relative power was computed as the ratio of the band power to the total broadband power in 1–250 Hz. For each electrode, the absolute and relative powers were averaged across all valid background windows of a given recording. The final output consisted of one row per mouse, recording day, and electrode, including the number of background markers used for averaging. For each segment: absolute power ($\mu V^2$): integral of the PSD in the band (bandpower); relative power (unitless): band power divided by total (1–250 Hz) power of the same segment and mean spectral density ($\mu V^2 \cdot Hz^{-1}$): absolute band power divided by band width (used in complementary analyses).

Longitudinal dynamics and slope comparison

For each region × band × cohort, a group-level linear model (OLS) was fitted,

$$Power_i = \beta_0 + \beta_1 day_i + \mathcal{E}_i,$$

with cluster-robust standard errors by mouse. We report slope, intercept, $R^2$, p-value, standard error, and n. Plots display day-wise means ± SEM and OLS predictions ± 95% CI. Slope differences between cohorts were tested via an interaction model,

$$power \sim day \times cohort_{Condition},$$

again, with mouse-clustered SEs. The interaction term (day:cohort) yields the slope difference (Ctrl–IHK) and its p-value.

Weekly distribution and between-cohort tests

For each electrode × band × week, we show boxplots of the per-mouse distribution (one value per mouse = median across days). Between-cohort comparisons (epileptic vs control) used two-sided Mann–Whitney U tests. We report raw p-values and q-values after Benjamini–Hochberg FDR, applied per band across weeks for that electrode; rank-biserial effect size.

$$r_{rb} = \frac{2U - n_1 n_2}{n_1 n_2}$$

For each cohort × electrode × band, we compared week pairs (W1↔W2, W2↔W3, W3↔W4, W1↔W4) using paired Wilcoxon tests on mice present at both weeks. We report raw p, FDR q-values per band (across the tested week pairs for that electrode/cohort), and effect size $r = z/\sqrt{n}$

Region-wise ANCOVA (cohort effect with adjustment)

For each region × band, we tested the cohort effect on power (here in relative units for the main run) with:

$$\text{power} \sim \text{cohort}_{\text{condition}} + \text{total}^{\text{dens}}_{1\text{–}250}(z) + \text{day}_c,$$

where $\text{total}^{\text{dens}}(z)$ is the z-score of 1–250 Hz mean spectral density computed at the mouse × day level (median) and $\text{day}_c$ is day centered (median). Standard errors were clustered by mouse. We report the cohort coefficient, 95% CI, p-value, and $R^2$. As a robustness check, a residualized version was also performed: (i) fit power $\sim \text{total}_z + \text{day}_c$; (ii) test cohort on the residuals.

Latency to the first seizure:

The latency to the first seizure was defined as the earliest day on which an iPD (ictal paroxysmal discharge) event was observed on a given electrode. If no iPD was detected on that electrode during the recording period, the mouse was considered censored at its last recorded day for that electrode. For both EF and CH electrodes, we constructed a survival dataset with one entry per mouse. Each entry contained: the time variable, defined as the first day of iPD occurrence (if observed) or the last recorded day (if censored), and the event indicator (1 = iPD observed, 0 = censored). This yielded electrode-specific survival tables describing the latency distribution across animals. Kaplan–Meier survival curves were fitted for each electrode using the lifelines Python package. Curves represent the probability of remaining seizure-free as a function of days post-KA. Confidence intervals (95%) were estimated, and censoring events were marked along the survival curves. Differences in seizure latency distributions between electrodes were tested using the log-rank test. Kaplan–Meier curves were plotted using matplotlib (Python 3.11)

Event construction, spike /HFOS rates and centrality (predictive analyses)

Events were grouped by mouse×day×electrode (and by region or hemisphere for aggregates). For each grouping we computed rates per minute as (count)/(recorded minutes).

We also computed hemispheric aggregates (mean rate across all regions of a hemisphere), hemisphere-only aggregates excluding the target region, and global laterality indices (LI):

$$\text{LI} = \frac{\text{ipsi} - \text{contra}}{\text{ipsi} + \text{contra} + 10^{-9}}.$$

A simple time-coincidence–based node strength (centrality) for the target region was computed during W1 (non-seizure windows) by counting spike co-occurrences within ±200 ms (tolerance ±5 ms) across regions, normalized by $\sqrt{n_a n_b}$

For each target electrode (EF, CH) and mouse we built a feature vector combining:

Electrode-local W1: spikes·$min^{-1}$, FR·$min^{-1}$, %FR (= FR/(FR+RP)), and (if available) relative PSD (relative) bands (delta/theta/beta/gamma/ripple).

Hemispheric aggregates (same hemisphere as the target): spikes/min and FR/min; and the same excluding the target region e.g. averaging all regions of the hemisphere except the target electrode's anatomical region to isolates hemispheric context outside the local node to avoid circularity.

Global laterality indices: LI spikes, LI FR, if positive: ipsilateral dominance. Permit the observation of how strongly the epileptic hemisphere dominates. High |LI| suggests lateralized pathology.

Target-region centrality: For the target region, we count time-coincident spikes with all other regions within ±200 ms (tolerance ±5 ms), normalized by $\sqrt{n_a n_b}$ to identify functional "hubness" by synchrony; higher implies the node sits in a strongly co-active network.

Hemispheric PSD (relative) Hemisphere-level relative power in a band, sometimes excluding the homologous DG to test whether prediction is driven by diffuse hemispheric content vs a single region. All features were calculated from non-ictal segments in W1.

For each mouse×electrode we computed a seizure rate over W4 (events per minute). Mice with strictly zero rate formed the "low" class. Among mice with non-zero rate, we thresholded at the median to define a binary label ($y = 1$ = higher-seizure; $y = 0$= lower/none). Labels were defined per electrode.

Leave-one-subject-out (LOSO) scoring and inference

Because each feature is a single scalar per mouse, LOSO reduces to evaluating the held-out mouse's feature as a score and computing the AUC from the set of (score, label) pairs across mice. For each feature×electrode we reported: LOSO AUC (primary metric), 95%. bootstrap CI from 10,000 mouse-level resamples, a permutation p-value for $H_1$ :AUC $>$0.5 (labels permuted 5,000 times; same scores), the direction (sign of $\bar{s}_{y=1} - \bar{s}_{y=0}$).

Multiple-testing control for prediction

To respect the heterogeneity of feature types, p-values were corrected within feature families and per electrode: Benjamini–Hochberg (BH) FDR → $q_{\mathrm{BH}}$, Storey's q-values with $\pi_0$estimated over $\lambda \in [0.05, 0.95]$→ $q_{\mathrm{Storey}}$. Features categories were: Lateralization, PSD, Spikes, HFO (FR), Centrality, and Other. Comparisons required ≥3 mice per group/condition (after week mapping and aggregation). Paired tests required mice with data in both weeks. All tests were two-sided unless specified (AUC permutation was one-sided: AUC>0.5).

Anchor-based co-spiking analysis

To quantify short-latency co-spiking relative to a hippocampal anchor (EF or CH), for each mouse–day we computed the probability that at least one spike in a target region occurred within a post-anchor window following each anchor spike:

$$P_{real}(\omega) = \frac{1}{N_{anchor}} \sum_{k=1}^{N_{anchor}} 1\{\exists\ target\ spike\ in\ [t_k + \omega_{start}\ , t_k +\ \omega_{end}]\}$$

The analysis used a single window [0,20]ms. A baseline was obtained by uniformly jittering anchor spike times by ±50 ms (clipped to the recording interval) and recomputing the probability over S = 200 shuffles; the baseline is the shuffle mean:

$$P_{\text{base}}(\omega) = \frac{1}{S} \sum_{s=1}^{S} P_{\text{jitter}}^{(s)}(\omega), S = 200.$$

The co-spiking excess probability was defined as

$$\Delta P(\omega) = P_{\text{real}}(\omega) - P_{\text{base}}(\omega).$$

We computed $\Delta P$for all anchor–target pairs per mouse–day, then collapsed to mouse means (mouse = unit of inference). Analyses were run separately with EF and CH as anchors. Two one-sided permutation tests (10,000 label flips) were used with Benjamini–Hochberg FDR across regions:

Sign test vs 0 (within period): does mean $\Delta P(w)$across mice exceed 0? $H_0$: $\mathbb{E}[\Delta P] = 0, H_1$: $\mathbb{E}[\Delta P] > 0.$

Paired W4 > W1 (within region): is $\Delta P(w)$larger in W4 than W1 for the same mice? $H_0$: $\mathbb{E}[\Delta P_{\text{W4}} - \Delta P_{\text{W1}}] = 0, H_1$: $> 0.$

For visualization, 2,000 bootstrap resamples over mice provided 95% CIs around the mean $\Delta P$.

HFO-conditionned co-spiking contrast:

To assess whether FR-tagged anchor spikes are more co-spike-effective than other spikes, we contrasted anchor subsets (e.g., FR vs none, FR vs RP). For each mouse–day, target region, and the [0,20]ms window:

$$\Delta P_{g_1-g_2}(w) = P_{\text{real}}^{(g_1)}(w) - P_{\text{real}}^{(g_2)}(w), \qquad \mathrm{g1, g2,} \in \{\mathrm{FR, RP, none}\}.$$

Values were collapsed to mouse means and submitted to the same permutation tests (sign vs 0; paired W4 > W1) with FDR across regions.

Spike time tiling coefficient (STTC) connectivity matrices

Pairwise synchrony independent of firing rate was estimated with the STTC using a coincidence tolerance $\tau = 35$ms and a minimum of 20 spikes per region per recording. STTC was computed for each mouse–day and then aggregated within period (W1, W4) by unbiased averaging (sum of valid values divided by their count), yielding matrices $M_{\mathrm{W1}}$and $M_{\mathrm{W4}}$(diagonals set to 0). A difference matrix $\Delta = M_{\mathrm{W4}} - M_{\mathrm{W1}}$was also reported. Heatmaps were displayed with color limits set to the 5th–95th percentiles of observed values.

Synchrony lateralization index (LI)

From each STTC matrix we computed a lateralization index contrasting within-hemisphere vs cross-hemisphere synchrony:

$$LI_{sync} = \frac{\overline{STTC}_{within-hemi} - \overline{STTC}_{cross-hemi}}{\overline{STTC}_{within-hemi} + \overline{STTC}_{cross-hemi} + \varepsilon},$$

where hemisphere labels were taken from the region map and $\varepsilon = 10^{-12}$prevents division-by-zero. Positive values indicate stronger within-hemisphere than cross-hemisphere synchrony.

Relative spike-timing order across regions (Kendall's W):

To test for a consistent temporal ordering of target regions relative to the anchor, we extracted, for each mouse–day, the nearest-neighbor lag (ms) of each target spike to anchor spikes within ±100 ms; days with ≥10 valid lags per region were retained. For each mouse, regional lags were summarized by the median, yielding a mouse × region matrix. On the core set of regions present in ≥3 mice, each mouse's lags were converted to ranks (average-tied), and Kendall's W (0–1) was computed separately in W1 and W4. 2,000 mouse-level bootstrap resamples provided 95% CIs.

Community structure and modularity

Graphs were built from STTC matrices (edge weight = STTC, non-positive weights excluded). For analysis, edges were thresholded at the 0.50 quantile of positive weights to avoid trivial single-community partitions. Community detection preferentially used Louvain (either python-louvain or the NetworkX implementation), with fallback to greedy modularity. We estimated modularity $Q$ in W1 and W4 and tested $\Delta Q = Q_{\mathrm{W4}} - Q_{\mathrm{W1}}$using node-label permutations where the same permutation was applied to W1 and W4 to preserve a coherent null for differences (5,000 permutations by default in code; one-sided). We also tested for a bi-hippocampal module at W4 (a community containing EF and CH) and reported its null frequency under label permutations. For visualization, we drew graphs using only edges above the 0.70 quantile of positive weights (clarity), colored nodes by community, and used node border color to indicate hemisphere.

Bi-hippocampal index (homotopic pairs)

We defined a bi-hippocampal index as the mean STTC across homotopic hippocampal pairs:

$$biHPC = mean\left\{STTC_{DG_{ipsi},DG_{contra}}, STTC_{CA1_{ipsi},CA1_{contra}}, STTC_{Sub_{ipsi},Sub_{contra}}\right\}$$

One-sided p-values were obtained by label permutations separately for W1 and W4, and Δ(W4–W1) was tested under a coherent null where the same permutation was applied to W1 and W4.

Unless otherwise stated, tests are one-sided (directed hypotheses: excess co-spiking > 0; W4 > W1) and p-values are FDR-adjusted (Benjamini–Hochberg) across the family of regions in each analysis. Bars/bands indicate mouse-level 95% bootstrap CIs. Heatmap limits use the 5th–95th percentiles of the data.

Directional cross-correlation between ipsilateral and contralateral dentate gyrus

Spike times were extracted from the same curated spike dataset used for all hippocampal analyses (see main Methods). For the present analysis, we restricted spikes to KA-treated animals (IHK condition) and to the two electrodes located in the ipsilateral (epileptogenic) and contralateral dentate gyrus (EF and CH, respectively; electrode IDs 11 and 6 in the chronic implant). Spike times were originally stored as sample indices at 2 kHz and converted to seconds for analysis.

For each spike, we assigned an event type by combining the regional and sub-event labels used throughout the study (sub_event_type when available, otherwise reg_event_type). Spikes belonging to ictal or peri-ictal events (classified as PD, iPD, sPD, or iPD_long in the main pipeline) were flagged and excluded from all analyses below, so that cross-correlation, Granger causality and transfer entropy were computed on interictal activity only.

To capture the time course of epileptogenesis, days post-KA were grouped into four latent periods: days 3–7 (W1), 9–13 (W2), 15–19 (W3) and 21–29 (W4). For each mouse and week, we then constructed two spike trains: all interictal spikes recorded on EF and all interictal spikes recorded on CH.

Spike-triggered cross-correlograms between EF and CH were computed using EF spikes as reference and CH spikes as target. For each reference spike time $t_{\mathrm{EF}}$, we collected all CH spikes within a symmetric time window $[t_{\mathrm{EF}} - \Delta, t_{\mathrm{EF}} + \Delta]$(typically $\Delta = 100$–$200$ ms), and accumulated the temporal differences $\Delta t = t_{\mathrm{CH}} - t_{\mathrm{EF}}$into a histogram with fixed-width bins (5 ms). This yielded, for each mouse–week pair, a cross-correlogram $C(\Delta t)$expressed as counts per lag bin. To account for differences in spike count across recordings, correlograms were normalized by the number of reference spikes, resulting in an estimate of the probability of observing a CH spike in each lag bin conditional on an EF spike.

For population-level visualization, normalized cross-correlograms were computed for each mouse–week and then averaged across mice within a given week. The mean and standard error of the mean were used to display the group-level cross-correlation profiles.

To quantify the directional balance of short-latency interactions, we defined an asymmetry index based on the integrated cross-correlation in a ±50 ms window around zero lag. For each mouse–week, we integrated the normalized cross-correlation separately over negative lags (−50 to 0 ms) and positive lags (0 to +50 ms), and computed an asymmetry index

$$A = \frac{\mathrm{area}_{\mathrm{pos}} - \mathrm{area}_{\mathrm{neg}}}{\mathrm{area}_{\mathrm{pos}} + \mathrm{area}_{\mathrm{neg}} + \varepsilon},$$

where $\text{area}_{\text{pos}}$ and $\text{area}_{\text{neg}}$ denote the integrated probability mass over positive and negative lags, respectively, and $\varepsilon$ is a small constant to avoid division by zero. Negative values of $A$ indicate that CH spikes tend to precede EF spikes (dominant CH→EF contribution), whereas positive values indicate the reverse.

In addition, we computed an effective lag for each mouse–week as the center-of-mass of the cross-correlation within ±50 ms,

$$\text{lag}_{\text{COM}} = \frac{\sum_{|\Delta t| \leq 50 \text{ ms}} \Delta t C(\Delta t)}{\sum_{|\Delta t| \leq 50 \text{ ms}} C(\Delta t)},$$

providing a single signed latency in milliseconds (negative values indicating that CH precedes EF on average). Both the asymmetry index and effective lag were used as summary measures of directional balance and were subjected to statistical analyses across weeks as described in Supplementary Results.

Granger causality on binned spike trains

To probe directed interactions at a slower timescale, we applied Granger causality (GC) to binned spike-count time series derived from EF and CH. For each mouse–week, interictal spikes from EF and CH were binned into non-overlapping time bins of fixed width (typically 10–20 ms), starting at 0 s and truncated to the largest multiple of a 1 s window fully contained in the recording. This yielded two integer-valued time series representing spike counts per bin in each region.

We segmented the binned spike-count series into consecutive 1 s windows (each consisting of a fixed number of bins). Windows with insufficient activity (total spike count < 3 in either region) were discarded to avoid numerical issues. Within each retained window, we assembled a two-dimensional time series

$$X(t) = [x_{\text{EF}}(t), x_{\text{CH}}(t)],$$

and standardized each component to zero mean and unit variance (z-score) to improve model stability.

Granger causality was estimated using standard vector autoregressive models as implemented in the *grangercausalitytests* function from the statsmodels library. For each direction (EF→CH and CH→EF) and each candidate lag order (up to a maximum lag, e.g. 10 bins), we compared the residual sum of squares of a full model including past values from both regions to that of a reduced model including only the past of the target region, yielding an F-statistic for the null hypothesis “past activity in the source region does not improve prediction of the target”. For each window and direction, we retained the maximum F-statistic across lags as a summary of GC strength in that direction. Mean GC values were then calculated across all windows for each mouse–week and direction, and a directional difference

$$\Delta\mathrm{GC} = \mathrm{GC}_{\mathrm{EF}\to\mathrm{CH}} - \mathrm{GC}_{\mathrm{CH}\to\mathrm{EF}}$$

was derived to characterize the relative causal influence of EF on CH versus the reverse. These per-mouse per-week values were used for group-level comparisons and regression across weeks (see Supplementary Results).

Transfer entropy between EF and CH

As a complementary, model-free measure of directed information flow, we computed transfer entropy (TE) between EF and CH using binarized spike trains. For each mouse–week, the same interictal spike times used for the GC analysis were binned into fixed-width bins (typically 20 ms), and each bin was converted to a binary value indicating the presence (1) or absence (0) of at least one spike in that bin. This yielded two binary time series representing spike occurrence in each region.

The binary sequences were segmented into consecutive 1 s windows, and windows in which one of the series was completely flat (all zeros) were excluded. For each remaining window, we estimated the transfer entropy from EF to CH and from CH to EF using the *transfer_entropy* function from the pyinform library, with a history length $k$(embedding order) chosen to match the timescale of interest (e.g. $k = 3$bins). This provides, for each direction, a non-parametric estimate of the information contributed by the past of the source series to the future of the target series, beyond the information contained in the target’s own past.

For each mouse–week, TE values were averaged across all windows to obtain mean TE in each direction, and a directional difference

$$\Delta\mathrm{TE} = \mathrm{TE}_{\mathrm{EF}\to\mathrm{CH}} - \mathrm{TE}_{\mathrm{CH}\to\mathrm{EF}}$$

was computed. These $\Delta\mathrm{TE}$ values were used as summary metrics of directed information flow and analyzed across weeks using linear regression and paired non-parametric tests.

### Supplementary Results

#### Early focal emergence of HFO-coupled spikes during epileptogenesis

Quantitative modeling confirmed a strong monotonic rise in spike rate within the epileptic hippocampus (Fig. 2D): spikes per minute increased significantly in both DG and CA1 on the injected side (slope = 1.3640, $R^2$ = 0.8655, $p < 0.0001$), whereas saline controls showed no comparable change (slope = –0.0421, $R^2$ = 0.1297, p = 0.21). Spike amplitude increased progressively (slope = +8.63 μV·day$^{-1}$, $R^2$ = 0.94, $p < 0.0001$; Fig. 2E), while half-amplitude width remained stable (slope = –0.077 ms·day$^{-1}$, $R^2$ = 0.02, p = 0.62; Fig. 2F). HFO-coupled spikes showed distinct temporal trajectories (Fig. 2G): fast-ripple–associated spikes (250–500 Hz) increased most steeply (slope = 0.5114, $R^2$ = 0.89, $p < 0.0001$), followed by ripple-associated (80–250 Hz; slope = 0.2042, $R^2$ = 0.76, $p < 0.0001$) and non-HFO spikes (slope = 0.6476, $R^2$ = 0.82, $p < 0.0001$). Pairwise slope tests confirmed significant ordering (fast-ripple > ripple > noHFO; all $p < 0.0001$). By week 4 (Fig. 2H), fast-ripple–associated spikes represented 22.6 ± 3.4 % of all discharges (W4 > W1, p = 0.0323), while non-HFO spikes decreased to 69.9 ± 2.8 % (p = 0.0171).

These quantitative data confirm that during the latent phase, epileptiform activity evolves from sparse, irregular spiking to a dense and structured pattern dominated by HFO-nested events - especially fast ripples - reflecting the progressive consolidation of hyperexcitable hippocampal microcircuits.

#### Delayed contralateral engagement and biphasic dynamics

Quantitative modeling confirmed that the contralateral hippocampus followed a delayed and weaker trajectory compared to the ipsilateral focus (Fig. 2J). Spiking activity increased modestly over days (slope = 0.4898, $R^2$ = 0.5313, p = 0.0031), while saline controls remained stable (slope = 0.0152, $R^2$ = 0.0533, p = 0.43). Spike amplitude showed a trend toward increase (slope = +6.59 μV·day$^{-1}$, $R^2$ = 0.23, p = 0.08; Fig. 2K), and half-amplitude width decreased significantly (slope = –0.22 ms·day$^{-1}$, $R^2$ = 0.54, p = 0.0029; Fig. 2L), confirming progressive sharpening of spike morphology. HFO-coupled events exhibited specific temporal slopes (Fig. 2M): ripple-associated spikes increased most strongly (slope = 0.2070, $R^2$ = 0.6842, p = 0.0003), followed by spikes without HFOs (slope = 0.2026, $R^2$ = 0.333, p =

0.0003) and fast-ripple–associated spikes (slope = 0.0759, $R^2$ = 0.639, p = 0.0006). Pairwise comparisons confirmed significantly greater increases for ripple vs. fast-ripple spikes (p = 0.0028), while fast-ripple and noHFO slopes did not differ significantly (p = 0.13). By week 4 (Fig. 2N), ripple-associated spikes represented 22.9 ± 2.5 % of contralateral discharges (W4 > W1, p = 0.040), whereas fast-ripple events rose modestly (3.7 → 7.3 %; p = 0.048) and non-HFO spikes declined (81.7 → 69.8 %; p = 0.017). These quantitative results demonstrate that while both hemispheres become progressively synchronized, they follow distinct temporal and spectral trajectories - fast-ripple predominance ipsilaterally and ripple predominance contralaterally - reflecting asymmetric yet coordinated maturation of epileptogenic dynamics.

**Contralateral hippocampus maintain stable spectral organization throughout epileptogenesis**

Longitudinal PSD analyses revealed minimal spectral change in the CH. Week-wise comparisons at W1 and W4 showed no significant shifts across theta, beta, or gamma bands after BH-FDR correction. The only deviation was a transient W1 delta difference (q=0.0177, r_rb=−0.85), which did not persist at later weeks (Fig. 3A). All remaining between-cohort weekly comparisons were non-significant.

Day-wise linear regression further confirmed spectral stability across the 4-week period. Delta relative power demonstrated a shallow, non-significant increase (slope=+0.00115·$day^{-1}$, $R^2$=0.0102, p=0.311), and no interaction with cohort (p=0.982). Theta similarly showed no longitudinal change (p=0.727; interaction p=0.179), as did beta (p=0.418; interaction p=0.49) and gamma (p=0.129; interaction p=0.155) (Fig. 3C, D,G,H). No band exhibited a significant cohort×day interaction, indicating no epileptogenesis-related divergence over time.

Paired within-mouse Wilcoxon week-to-week comparisons (W1↔W2↔W3↔W4) yielded no significant effects after BH-FDR correction across all four frequency bands, demonstrating absence of progressive spectral remodeling. Together, these findings indicate that contralateral hippocampal background activity remains spectrally stable despite ongoing epileptogenic processes in the ipsilateral hemisphere.

**Epileptic focus undergoes progressive and structured spectral reorganization**

In contrast, the EF displayed robust and evolving spectral alterations (Fig. 3B). By W1, epileptic mice differed significantly from controls across all major frequency bands: delta decreased (q=0.009, r_rb=−0.82), while theta (q=0.0009, r_rb=−1), beta (q=0.00047, r_rb=−1), and gamma (q=0.00047, r_rb=−1) were elevated. These effects strengthened by W4, again spanning all bands - delta (q=0.0009, r_rb=−1), theta (q=0.003, r_rb=−0.88), beta (q=0.00062, r_rb=−0.97), and gamma (q=0.00047, r_rb=−0.97) - indicating widening divergence from controls.

Day-wise modeling revealed structured, monotonic remodeling across epileptogenesis. Delta relative power steadily declined (slope=$-0.0063 \cdot day^{-1}$, $R^2$=0.204, p=$2.49 \times 10^{-7}$), whereas theta increased (slope=$+0.00333 \cdot day^{-1}$, $R^2$=0.18, p=0.0007) and beta increased (slope=$+0.00149 \cdot day^{-1}$, $R^2$=0.118, p=0.0087). Gamma showed a weaker, non-significant upward trend (slope=$+0.0009 \cdot day^{-1}$, $R^2$=0.0302, p=0.156). Significant group×day interactions confirmed epileptogenesis-specific divergence for delta (p=0.0003), theta (p=0.003), and beta (p=0.014), but not gamma (p=0.187) (Fig. 3E–J).

Paired within-mouse W1→W4 analyses validated progressive spectral redistribution. Delta power decreased (W=2, q=0.0029, r=−0.94), while theta (W=7, q=0.019, r=0.79), beta (W=4, q=0.0068, r=0.87), and gamma (W=11, q=0.027, r=0.685) increased significantly. No contralateral comparison survived the same correction threshold, confirming hemispheric specificity. Although EF spectra diverged strongly from controls early in epileptogenesis, the subsequent redistribution of spectral power (delta↓, theta/beta↑) progressively realigned the EF oscillatory profile toward control-like proportions, such that by the end of the monitoring period, IHK mice displayed a background spectral composition more similar to controls than at W1, reflecting partial re-equilibration rather than continuous deterioration. Collectively, these results demonstrate that epileptogenesis is accompanied by a gradual, coordinated shift in background activity at the seizure focus - characterized by delta suppression and increased theta–beta power - emerging early, progressing over weeks, and remaining anatomically restricted to the ipsilateral hippocampus.

**Evolution, amplification and predictive structure of paroxysmal discharges**

To determine whether epileptogenesis accelerates seizure expression in the epileptic hemisphere, we compared time to first iPD using Kaplan–Meier survival estimates. EF and CH showed nearly identical survival curves, with overlapping 95% confidence intervals (Fig. 4A). A log-rank comparison confirmed no hemispheric difference in seizure latency ($\chi^2$ = 0.018, p = 0.894; $-\log_2$ p = 0.162), indicating that initial ictal emergence is bilaterally symmetric despite unilateral KA administration. Mice without seizures were treated as right-censored at their last recorded day.

We next tested whether early epileptiform activity predicts subsequent seizure severity. Week1 spike rate was not correlated with week4 seizure rate in either hemisphere (CH: Spearman ρ = −0.52, p = 0.070; EF: ρ = 0.02, p = 0.957; Fig. 4B–C). Thus, early spiking frequency alone is not a reliable biomarker of emerging ictogenesis.

Longitudinal modeling of dentate iPDs revealed distinct hemispheric trajectories (Fig. 4D–K). In CH, iPD rate showed a non-significant upward trend (slope = +0.0068 $min^{-1}\cdot day^{-1}$, $R^2$ = 0.12, p = 0.092), whereas iPD duration increased significantly over time (+0.73 $s\cdot day^{-1}$, $R^2$ = 0.19, p = 0.0084). ISI increased modestly (+6.45 $ms\cdot day^{-1}$, $R^2$ = 0.13, p = 0.086), and amplitude showed a near-significant rise (+19.8 $\mu V\cdot day^{-1}$, $R^2$ = 0.24, p = 0.053). In contrast, EF demonstrated stronger and more consistent escalation across all features: iPD rate increased robustly (+0.0095 $min^{-1}\cdot day^{-1}$, $R^2$ = 0.16, p = $3.0\times10^{-4}$), duration rose steadily (+0.62 $s\cdot day^{-1}$, $R^2$ = 0.10, p = 0.026), ISI shortened markedly (−9.97 $ms\cdot day^{-1}$, $R^2$ = 0.18, p = $2.0\times10^{-4}$), and amplitude increased (+9.24 $\mu V\cdot day^{-1}$, $R^2$ = 0.11, p = 0.011). Standard errors were clustered by mouse. These trends indicate progressive temporal densification and recruitment within the epileptic dentate gyrus.

We further compared regionally isolated vs. bilateral hippocampal PDs (Bi-HPC) across hemispheres (Fig. 4L–O). In CH, duration decreased in isolated events (slope = −0.108 $s\cdot day^{-1}$, $R^2$ = 0.54, p = 0.0027) and trended downward in Bi-HPC discharges (−0.172 $s\cdot day^{-1}$, $R^2$ = 0.28, p = 0.054), with a significant between-group difference (p = 0.0011). During the chronic stage (days 21–29), Bi-HPC events lasted longer (17.81 s) than isolated ones (5.93 s; Mann–Whitney p < 0.0001, r = 0.38). Amplitudes increased similarly across both CH event types (isolated +8.68 μV·day$^{-1}$, $R^2$ = 0.50, p = 0.0050; Bi-HPC +12.37 μV·day$^{-1}$, $R^2$ = 0.54, p = 0.0029), with no significant difference (p = 0.38).

In EF, both event classes lengthened significantly, but Bi-HPC events showed markedly steeper duration growth (+0.646 s·day$^{-1}$, $R^2$ = 0.73, p = 0.0001) compared to isolated ones (+0.115 s·day$^{-1}$, $R^2$ = 0.57, p = 0.0017), with a highly significant between-group difference (p < 0.0001). During the chronic period, Bi-HPC events were substantially longer (24.82 s vs. 11.46 s; p < 0.0001, r = 0.33). Amplitude increased sharply in both isolated (+7.93 μV·day$^{-1}$, $R^2$ = 0.93, p < 0.0001) and Bi-HPC events (+9.12 μV·day$^{-1}$, $R^2$ = 0.82, p < 0.0001). Between-group slope differences were non-significant (p = 0.39).

Finally, we evaluated whether early electrophysiological features could classify week4 seizure burden using LOSO machine-learning prediction. In CH, the strongest predictor was global spike lateralization (AUC ≈ 0.74; p = 0.092; q = 0.184), though not surviving FDR correction. All other features performed near chance. In EF, global fast-ripple lateralization yielded the highest performance (AUC ≈ 0.85; p = 0.037; q = 0.075), representing a trend after FDR adjustment. Absolute W1 spike or FR rates alone showed poor discriminability (AUC ≤ 0.62). Thus, early hemispheric asymmetry - rather than overall event load - may serve as a more informative prognostic marker of seizure severity.

**Progressive strengthening and bilateral reorganization of co-spiking networks**

Short-latency co-spiking probability (ΔP; Fig. 5A–B) was significantly elevated above baseline during the first week post-SE across multiple regions (all p_FDR < 0.05), including contralateral PrL, Cg1, M2, and V1, as well as ipsilateral Cg1, PrL, Ment, Sub, and CA1_contra. This widespread synchrony indicates rapid bilateral propagation of EF spikes during early epileptogenesis. By week 4, ΔP remained significantly elevated in multiple ipsilateral regions and extended to subicular, prefrontal, motor, and visual cortices. Within-mouse comparisons (W4 > W1) revealed a significant increase in V1_ipsi (p_FDR = 0.037), with stable or upward trends in other regions.

For fast-ripple–conditioned analyses (Fig. 5C), within-week contrasts between FR-tagged and non-HFO ("no-HFO") spikes yielded no significant differences after FDR correction at W1 or W4 (all p_FDR > 0.05). However, longitudinal comparisons (W4 > W1) revealed significant positive shifts in M2_ipsi (p_FDR = 0.003), Cg1_contra (p_FDR = 0.0067), PrL_contra (p_FDR = 0.0115), CA1_contra (p_FDR = 0.0135), and Cg1_ipsi (p_FDR = 0.047).

At W1, mean ΔP_FR–none values were slightly negative (–0.011 ± 0.004), suggesting that early FR events did not efficiently drive remote synchrony, consistent with resonance in localized epileptogenic microcircuits. By W4, ΔP_FR–none approached zero or became positive in several regions (+0.009 ± 0.005), reflecting loss of FR confinement and broader network entrainment.

Together, these data demonstrate a progressive strengthening of rapid co-spiking across hemispheres, with early, widespread fronto-limbic synchrony gradually incorporating sensory and contralateral nodes. The transition from locally confined to bilaterally integrated fast-ripple coupling marks the consolidation of a distributed epileptic network.

**From focal synchrony to bilateral integration of mesoscale networks**

Pairwise STTC analyses (τ = 35 ms; Fig. 5D–F) revealed significant global strengthening of short-latency synchrony between W1 and W4. The largest increases occurred along the hippocampo-subicular pathway (DG–Sub: ΔSTTC = +0.084 ± 0.019, p = 0.0004; Sub–CA1: +0.067 ± 0.015, p = 0.0009) and across hemispheres (EF–CH: +0.052 ± 0.013, p = 0.0011). Long-range cortico-cortical couplings (e.g., PrL–Cg1, M2–V1) showed minimal change (p > 0.2), underscoring the selective reinforcement of hippocampal synchrony.

Graph-based topology of the strongest 25 % of STTC edges (Fig. 5G–H) confirmed a reorganization toward a more integrated structure. The bi-hippocampal index increased from 0.117 ± 0.008 to 0.193 ± 0.012 (p = 0.0068), and modularity (Q) rose from 0.044 ± 0.003 to 0.056 ± 0.004 (p = 0.0002). The lateralization index remained slightly negative (–0.199 → –0.221), indicating bilateral strengthening without hemispheric dominance.

To assess temporal ordering, we computed each region's median spike lag relative to EF within ±100 ms. Cross-animal rank concordance (Kendall's W) decreased from 0.25 (W1) to 0.14 (W4), demonstrating reduced stereotypy of recruitment despite stronger global coupling.

Both ΔP and STTC metrics showed a monotonic increase in inter-hippocampal synchrony across animals (Kendall's W = 0.72, p < 0.001). These results indicate that epileptogenesis follows a structured progression from early, distributed cortical engagement to the emergence of a stable, self-sustaining bi-hippocampal core - signifying the maturation of the epileptic network.

**Directional coupling and causal interactions during epileptogenesis**

To assess whether the progressive bilateral coactivation between EF and CH was accompanied by a change in directionality, we quantified the temporal asymmetry and effective lag of their spike cross-correlations across weeks. Cross-correlograms displayed a sharp central peak already present during the first week, indicative of near-synchronous activation of both dentate gyri.

The asymmetry index, defined as the normalized difference between positive and negative lag windows (0–50 ms vs −50–0 ms), was significantly negative during the early latent phase, indicating a slight precedence of CH spikes over EF spikes. This index gradually increased toward zero across weeks (linear regression: $p = 0.055$; paired Wilcoxon W1 < W4, $p = 0.02$), consistent with a progressive equalization of short-latency coactivation.

The effective lag (center of mass within ±50 ms) remained close to zero throughout the latent period (median ≈ −2 ms, $p = 0.41$), confirming a stable and nearly synchronous coupling between both regions. Directional Granger causality (ΔGC = EF→CH − CH→EF) and transfer entropy (ΔTE = EF→CH − CH→EF) computed on binned spike trains (10–20 ms bins, 1 s windows) showed no significant direction-specific trend across weeks ($p > 0.2$).

Together, these analyses indicate that hippocampal epileptogenesis does not entail a progressive dominance of one direction over the other, but rather a gradual symmetrization of bidirectional coupling between the ipsilateral and contralateral dentate gyri.

**Quantitative analysis of chemogenetic silencing effects during the latent phase**

Linear regression analysis confirmed a comparable increase in EF spike rate between groups (IHK: slope = 1.3640, $R^2 = 0.8655$, $p < 0.0001$; DFIHK: slope = 1.7633, $R^2 = 0.9195$, $p < 0.0001$; difference n.s.; Fig. 8A). Spike amplitude also rose similarly (IHK: slope = 8.6305, $R^2 = 0.9363$; DFIHK: slope = 9.3429, $R^2 = 0.9202$; both $p < 0.0001$; Fig. 7B).

Spike subtype distributions (Fig. 8C) evolved similarly in both groups, shifting from predominantly noHFO to sRP and sFR events. Quantitatively, the rate of noHFO spikes increased more steeply in silenced animals (IHK: 0.6476, $R^2$ = 0.8197 vs. DFIHK: 1.0303, $R^2$ = 0.9070; p = 0.0031; Fig. 8D). RP-associated spikes showed a similar pattern (IHK: 0.2042, $R^2$ = 0.7622 vs. DFIHK: 0.3114, $R^2$ = 0.9232; p = 0.0105; Fig. 8E). FR-associated spikes followed parallel trajectories (IHK: 0.5114, $R^2$ = 0.8915 vs. DFIHK: 0.4198, $R^2$ = 0.8577; p = 0.1993; Fig. 8F).

In the contralateral hippocampus (Fig. 8G–H), spike rate increased in both groups (IHK: 0.4898, $R^2$ = 0.5313, p = 0.0031; DFIHK: 0.6791, $R^2$ = 0.8976, $p < 0.0001$), with no significant group difference. Spike amplitude slopes were low and nonsignificant (IHK: 6.5876, $R^2$ = 0.2348, p = 0.0791; DFIHK: 1.7875, $R^2$ = 0.0337, p = 0.5297).

Subtype analysis in CH (Fig. 8I–L) showed that noHFO spikes increased significantly faster in silenced animals (IHK: 0.2026, $R^2$ = 0.3330 vs. DFIHK: 0.3973, $R^2$ = 0.8980; p = 0.0330; Fig. 8J). RP-associated spike slopes did not differ (IHK: 0.2070, $R^2$ = 0.6842 vs. DFIHK: 0.1724, $R^2$ = 0.8072; p = 0.4641; Fig. 8K), and FR-associated spike slopes were likewise similar (IHK: 0.0759, $R^2$ = 0.6388 vs. DFIHK: 0.1095, $R^2$ = 0.8791; p = 0.0963; Fig. 8L).

Overall, chemogenetic inhibition of the EF during the latent phase fails to prevent the progressive rise in epileptiform spiking. The modest enhancement of non–HFO-coupled discharges in silenced animals may reflect a compensatory adjustment in network excitability rather than genuine suppression of epileptogenesis.

**Quantitative analysis of contralateral silencing effects during the latent phase**

In the CH, spike rate increased similarly across groups (Fig. 9A), with slopes of 0.4898 $spikes \cdot min^{-1} \cdot day^{-1}$ ($R^2$ = 0.5313, p = 0.0031) for KA-treated animals and 0.5045 $spikes \cdot min^{-1} \cdot day^{-1}$ ($R^2$ = 0.5193, p = 0.0036) for contralaterally silenced mice. Spike amplitudes also followed parallel trajectories (Fig. 9B; IHK = 6.5876 $\mu V \cdot day^{-1}$, $R^2$ = 0.2348, p = 0.0791; CH-silenced = 6.3312 $\mu V \cdot day^{-1}$, $R^2$ = 0.2435, p = 0.0730).

Spike subtype proportions were stable across groups (Fig. 9C–F). NoHFO spike rates were comparable (IHK = 0.2026, $R^2$ = 0.3330, p = 0.0307; CH-silenced = 0.1934, $R^2$ = 0.3088, p = 0.0391; p = 0.9377; Fig. 9D). sRP also showed similar slopes (IHK = 0.2070, $R^2$ = 0.6842; CH-silenced = 0.2184, $R^2$ = 0.5886; p = 0.8649; Fig. 9E), as did sFR (IHK = 0.0759, $R^2$ = 0.6388; CH-silenced = 0.0992, $R^2$ = 0.5578; p = 0.4434; Fig. 9F).

In sharp contrast, pronounced effects emerged in the EF. Silenced animals exhibited a markedly steeper rise in spike rate (IHK = 1.3640, $R^2$ = 0.8655 vs. CH-silenced = 2.4668, $R^2$ = 0.9360; $p < 0.0001$; Fig. 9G) and significantly higher spike amplitudes (IHK = 8.6305 $\mu V \cdot day^{-1}$, $R^2$ = 0.9363 vs. CH-silenced = 11.4252 $\mu V \cdot day^{-1}$, $R^2$ = 0.9509; $p < 0.0001$; Fig. 9H).

Subtype-specific analyses revealed that noHFO spike rates increased similarly across groups (IHK = 0.6476, $R^2$ = 0.8197 vs. CH-silenced = 0.7135, $R^2$ = 0.6924; p = 0.6860; Fig. 9J), while sRP were significantly higher in silenced animals (IHK = 0.2042, $R^2$ = 0.7622 vs. CH-silenced = 0.3447, $R^2$ = 0.8934; p = 0.0032; Fig. 9K). sFR exhibited the most pronounced difference, with slopes nearly tripled in silenced animals (IHK = 0.5114, $R^2$ = 0.8915 vs. CH-silenced = 1.4016, $R^2$ = 0.9160; $p < 0.0001$; Fig. 9L).

These quantitative results confirm that silencing the contralateral hippocampus intensifies epileptiform dynamics in the epileptogenic focus. Rather than suppressing hyperexcitability, contralateral inhibition disrupts a putative interhemispheric balancing mechanism, thereby unmasking a latent facilitation of fast ripple–driven synchrony and accelerating the maturation of pathological network activity.
